# Recent Advancement in Biosensors Technology for Animal and Livestock Health Management

**DOI:** 10.1101/128504

**Authors:** Suresh Neethirajan, Sheng-Tung Huang, Satish K. Tuteja, David Kelton

## Abstract

The term *biosensors* encompasses devices that have the potential to quantify physiological, immunological and behavioural responses of livestock and multiple animal species. Novel biosensing methodologies offer highly specialised monitoring devices for the specific measurement of individual and multiple parameters covering an animal’s physiology as well as monitoring of an animal’s environment. These devices are not only highly specific and sensitive for the parameters being analysed, but they are also reliable and easy to use, and can accelerate the monitoring process. Novel biosensors in livestock management provide significant benefits and applications in disease detection and isolation, health monitoring and detection of reproductive cycles, as well as monitoring physiological wellbeing of the animal via analysis of the animal’s environment. With the development of integrated systems and the Internet of Things, the continuously monitoring devices are expected to become affordable. The data generated from integrated livestock monitoring is anticipated to assist farmers and the agricultural industry to improve animal productivity in the future. The data is expected to reduce the impact of the livestock industry on the environment, while at the same time driving the new wave towards the improvements of viable farming techniques. This review focusses on the emerging technological advancements in monitoring of livestock health for detailed, precise information on productivity, as well as physiology and well-being. Biosensors will contribute to the 4^th^ revolution in agriculture by incorporating innovative technologies into cost-effective diagnostic methods that can mitigate the potentially catastrophic effects of infectious outbreaks in farmed animals

## 1. Introduction

Advances in engineering research and biomaterials, coupled with the decreasing costs of electronic technologies, have resulted in the emergence of ‘sensing solutions’ and smart computing technologies that include internet and cloud–based connectivity to develop integrated and networked physical devices for data collection and analysis. These systems are equipped to automatically collect data on physiological parameters, farm environment, production measures and behavioural traits.

In the modern world, new diseases that threaten animals’ health emerge every year. There is currently a lack of reliable, cost-effective diagnostic tests for early detection of diseases in farmed livestock animals. Biosensing technologies have the potential to address these problems by developing innovative diagnostic tools for the rapid detection of key health threats within the agri-food livestock sector.

There are numerous factors that affect food production and have an influence on food security around the world. By 2050, food demand is expected to increase by 70%, and meat production will increase by 50%, making agri-food and livestock key industries for future growth (Alexandratos and Bruinsma, 2012). Health threats to animal populations can disrupt food supply chains and commerce with potentially long-lasting effects on human health, as well as economic impacts. With novel infectious agents and global pandemic factors on the rise in farmed livestock industries, efficient and timely strategies for monitoring and predicting risks are crucial. With current technology, detecting diseases in the early stage requires time-consuming and expensive laboratory tests. There is a need for detection tools that can predict when an incident is likely to occur and in what population, inform diagnosis and treatment options, and forecast potential impacts on a given population (both human and animal). Furthermore, such technologies must be accurate, affordable and broadly available. Strengthened laboratory and field capabilities are needed to support these capacities. Diagnostic tools provide crucial information to surveillance programs in diverse operational contexts, including networks and reference diagnostic laboratories associated with the World Organization for Animal Health (OIE), the United Nations Food and Agriculture Organization (FAO), and the Canadian Food Inspection Agency (CFIA). These systems not only integrate the data on individuals and groups; they can also help with the decision-making process by assisting in the early detection of health issues and wellbeing problems in individual animals. These integrated systems will also help in the implementation of corrective measures and improvements in management processes for animal husbandry practices. The biosensor market for the year 2013 was valued at US $11.39 Billion and is expected to increase to US$22.68 Billion by 2020. This growth in the biosensor market and associated applications is attributed to an increase in the demand for point-of-care testing. Furthermore, non-invasive health monitoring is also driving the growth and development of nanotechnology-based biosensors (Research, 2014). The precision farming market, important in livestock management, is expected to grow from USD 3.20 Billion in 2015 to USD 7.87 Billion by 2022 (MarketsandMarkets). The driving socio-economic and environmental factors that are expected to further the research and development to provide food for the growing human population are detailed in Figure 1.

**Figure 1.**
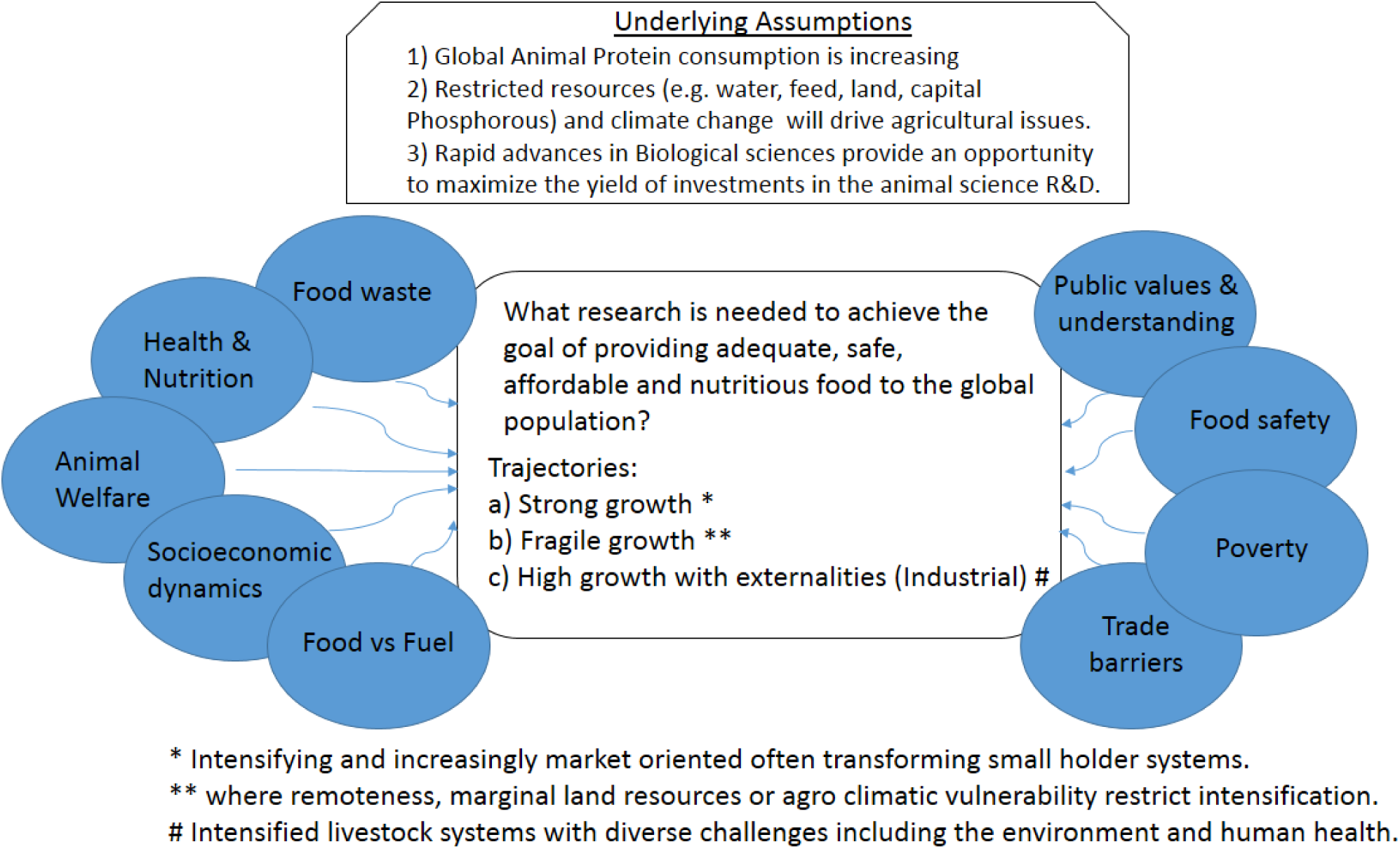
Schematic representation of environmental and socio-economic factors impacting future food production. Source: Force et al., (2015).

The market for point-of-care testing in veterinary diagnostics is expected to increase at a compound annual growth rate (CAGR) of 18%, reaching US$6.71 Billion by 2021. Novel diagnostic tools and disease modelling will enable decision-making and investigate the rapid diagnosis of epidemic and emerging diseases of farmed animals. The nanotechnology approach in developing biosensing tools offers direct benefits through simpler testing, smaller size, greater accuracy, faster results, and faster responses to key health threats in the farm animal sector.

We have entered a ‘fourth revolution’ in agriculture. This denotes the proliferation of new technologies including the Internet of Things, precision agriculture and mobile apps for disease surveillance. This review study will covers the technologies that will improve the capacity and efficiency of novel infection diagnosis, providing information needed to formulate sustainable risk assessment-based infection control programs for agri-food and livestock producers.

In this review, the focus relies on the emergent bio-sensing technologies that have the ability to transform management in the livestock industry and the methods associated with it. One of the salient features of biological, agricultural and environmental applications of nanotechnology is that the nanoscale devices and systems are of the same size-scale as biomolecules. While conventional sensors have been used in livestock monitoring, and as tools to assist in health monitoring and disease diagnosis, nanobiosensors have the ability to multiplex the bioassays on-site, thereby eliminating the need for the transportation of biological samples to centralised laboratories for analysis. Integration of these sensors for wireless data transfer via a server or through cloud-based systems would enable access to the analysed data for any internet-enabled device. Nanobiosensor applications will not only reduce the incumbent costs for reagents, sample handling, analysis times and transportation costs, but will also help in adapting and promoting sustainable agricultural techniques and ethical handling of livestock.

These technologies shall focus on the non-invasive methodologies to assess animal welfare by quantifying the stress and metabolic disease biomarkers, welfare assessment based on activity of the animals (monitoring oestrus and lameness detection to maximise animal production) and sensors for temperature and pH sensing (to determine calving alert and rumen function).

Furthermore, various non-invasive sensing technologies for early disease detection shall help in saving animals’ lives as well as reducing expenses for the farmers. A combination of these technologies and the use of ‘smart’ husbandry support systems will ensure maximum productivity while improving the wellbeing of farm animals (Caja et al., 2016). Non-invasive technologies for the chemical and biological analysis of samples from livestock, food and feed can rapidly provide detailed information to evaluate the safety of various biological samples. Biosensors equipped with robust data collection and integration infrastructure will be able to realise this potential and shall become vital elements in real-time analysis of industrial agriculture. Not only will such systems help in maximising the utilisation of resources for farming; they will also allow for an evaluation of individual and group behaviour of animals. Development of on-site biosensing technologies will enable rapid, cost-effective and meaningful monitoring of dietary inputs, environmental conditions, genetic makeup, performance, metabolism, welfare and physiological state of animals.

### 1.1 Biosensing - Taking a systems biology approach

The application of biosensors in animal husbandry and agriculture will increase competitiveness in the ever-changing global economy. The enormous amount of data generated by the continuous monitoring shall generate new knowledge on animals’ health and physiology and is expected to result in the development of technologies that will improve efficiency of animal production, better usage of dietary resources, improved health and welfare of animals through improved animal management, and reduced output of waste per unit of food product, thereby decreasing the impact of animal production systems on the environment.

Taking a systems biology approach to animal productivity and wellbeing is the way forward in animal husbandry, as well as in agriculture. It will also be indicative of future human performance and wellbeing initiatives. Collecting individual animal data as opposed to just ‘herd management’ will be necessary to monitor the wellbeing of individual animals as well as animal groups, and will help in identifying diseased animals sooner so as to provide healthcare and prevent any disease outbreaks. Moving agriculture (animal productivity and wellbeing) on a parallel course with human medicine and social science (human productivity and wellbeing) will enable the determination of multi-parametric data on physiological and environmental factors affecting animal welfare and productivity. This integrated database can be used to implement best farm practices to ensure animal welfare and productivity and to predict animal behaviour. Biosensing applications for livestock management and welfare will foster productive, value-added partnerships in ways that will lead to social, health, environmental and economic benefits. The approach to develop new solutions ranges from involving molecules to ecosystems, from nanotechnology to big data analysis and management, and from microbes to sheep to human populations. Moreover, real-time monitoring of animal health and assessment will have a direct impact on animal productivity and better utilisation of resources. The future of biosensors lies in utilising the comprehensive knowledge of animal physiology, genetics, environmental sciences and animal nutrition, and integrating this knowledge in a meaningful way will aid in the translation into real commercial and societal benefits.

### 1.2 Integrated in “decision support systems”

Animals contribute to the human society in a number of different ways. These include being a source of food, acting as models for studying human metabolism and diseases, and providing companionship and services. However, considering that the human population is expected to surpass 9 billion by 2050, feeding the population represents a formidable challenge. The global demand for food is expected to increase by more than 50%, while the consumption of animal food and food products is expected to rise by 73% by the year 2050 (Alexandratos and Bruinsma, 2012). The increase in human population demands increased availability of food using sustainable methods and implementation of strategies to increase animal productivity and agricultural production. It also demands the strengthening of legal systems to ensure sustainable food chain management. In a society increasingly concerned with the welfare of animals, methods to ensure ethical practices for animal rearing and animal wellbeing need to be implemented.

As the global demand for high-quality animal protein increases, it will result in a competition for limited food and feed sources from plants, further pressuring the agricultural industry to increase productivity of food and feed crops per unit area of arable land. The environmental impacts of livestock production (e.g., on soil, water, atmosphere and forest reserves) are key challenges to bear in mind as we devise plans and policies to manage and further develop the production systems that are environmentally sustainable, economically viable, ethically acceptable, and provide wholesome and nutritious food for animals and humans on a global scale. These challenges undoubtedly demand effective, rapid application of our cumulative knowledge, and innovative technologies in managing and caring for animals. Moreover, strong scientific bases for environmental policies, assessment of food attributes, safety of animal foods, policies and regulations governing animal management and welfare guidelines must form an essential platform for devising future technologies. Integrating data and knowledge gained from decades of research in environmental sciences, animal husbandry, agricultural practices and animal nutrition and behaviour with the modern integrated electronic systems will play a pivotal role in the optimisation and management of animal health and wellbeing (e.g., precision livestock management) and to improve sustainability of the supply chain for feed and food production.

This integration will be handled by decision support systems, which, to be most effective, must be robust under varying conditions; include technologies for rapid (automated) data collection via wireless data transmission systems (Ruiz-Garcia et al., 2009) (i.e., animal and environmental sensors); have substantial computing capacity for data analyses; have systems optimised to inform decision making and be reasonably easy to operate. The next breakthrough will be for these systems to use ‘real-time biometry,’ functioning in real time to monitor and control genotype, environment, wellbeing, productivity and animal product quality.

Development of novel methods for the real-time assessment and management of animal productivity and wellbeing is essential for investigating how the health-related parameters are affected by diet, animal husbandry, environmental factors, and genotype by environmental interactions. One of the key aspects of managing an animal’s environment will be to optimise feed ingredients and food quality (i.e., the most critical environmental factor), while maximising the use of co-products. The application of these novel integrative technologies shall be of interest primarily to the small livestock (chickens, pigs, fish, sheep), as well as companion animals (cats and dogs).

The precisely controlled small-scale experimental investigations on these species, focussed on improved understanding of physiology, are likely to be scaled up to pilot-scale and eventually result in future commercial applications. As for the livestock species, the quality of final product like meat, milk, eggs, etc. will require careful consideration for quality, with the ultimate aim of commercialisation of these integrated systems to be adapted on a larger world-wide scale where data and inputs from animals are directly incorporated for use in the management software.

### 1.3 Biosensors will manifest themselves as indispensable tools in animal husbandry

#### 1.3.1 Innovation and development for new approaches

Future developments in biosensors are expected to result in the development of new methodological and technological approaches to measuring dynamic changes in real time, with respect to the changes in physiological state and metabolism (e.g., gastrointestinal flora, circulating levels of anabolic and catabolic hormones, immune function, gene expression). This is to better understand the factors influencing animals’ responses, and to develop solutions (e.g., husbandry practices, technology and associated decision support system) that improve productivity and/or wellbeing of these animals.

#### 1.3.2 Real-time data acquisition and analysis

Monitoring of real-time autonomic responses (e.g., respiration rate, heartrate and heartrate variability, blood pressure, changes in peripheral blood flow) and defence-related reflexes (e.g., startle) using novel biosensing tools will help to investigate how housing, diet and genotype affect animals’ resiliency to stressors. These sensors will help in the understanding of factors that influence the wellbeing of animals, and in the development of solutions (e.g., husbandry practices, genotype selection) that improve the welfare of livestock and companion animals. Advances in wearable or imprinted biosensors that are flexible and allow data transfer remotely will be of special significance in this advancing area (Neethirajan, 2017).

#### 1.3.3 Rapid characterisation of food and feed

Biosensors shall be used to develop approaches enabling the rapid, accurate characterisation of dietary inputs and final products (meat, eggs, milk) in terms of nutrient content (total and bioavailable), anti-nutritional factors and bioactive components, as well as chemical and microbiological contaminants, with the aim of implementing this technology at the level of the commercial feed mill or animal food product processing plant. On the other hand, they would also help in the decision-making process to alter the composition of feed to the animals in case the animal products deviate from the expected nutritional status.

#### 1.3.4 Animal trait analysis and selection of robust breeds

Biosensing may also help to select special animal breeds that are robust and resilient to environmental stressors by enabling rapid assessment of the impacts of animal genotype and environmental factors at different life stages. Such assessment would yield critical knowledge to better understand genotype by environment interactions, in order to improve production efficiency and animal wellbeing. The developments in biosensing will also help us better predict and manage the impacts of climate change on animal agriculture over the next several decades.

#### 1.3.5 Enabling planning of energy budgets and reduction of environmental impact

The data collected and analysed using biosensors can assist in constructing detailed nutrient, energy and elemental budgets for diverse livestock species at different life stages in response to modulation in diet composition and environmental conditions, allowing precise management and efficient usage of nutrients and minimisation of waste outputs. This will have a direct impact on the efficient management of feed inputs and water resources while reducing the cost of production, wastages and environmental footprint. For example, real-time monitoring of cattle movement can provide information on the quality and quantity of forage, and the ability to determine required changes to the grazing systems. Monitoring variables like the consumption of water can provide insights into feeding behaviour, as well as the interaction between grazing systems management and this behaviour. Quantifying animal water consumption within a grazing environment can help to identify the impact of animal grazing on water quality, as well as land utilisation. (Davis, 2007a; Davis, 2007b)

#### 1.3.6 Development of mathematical algorithms for better understanding of complex biological systems and their interaction with the environment

In the present day and age of big data, the data from animal farms is expected to help in the development of advanced bio-mathematical models that are able to integrate data from the aforementioned scientific research efforts and theoretical understanding of complex biological systems. These models and simulations will allow for an improved quantitative appreciation of the scientific and management aspects of animal agriculture. These will enable the assessment of changes in the system with respect to different production, genetic selection, nutritional and environmental factors. Ultimately, these models will help to identify approaches and strategies to improve the productivity, efficiency and wellbeing of animals and mitigate the potential negative environmental impacts of livestock production. These models will also provide the basis for the development of the specific algorithms required by a variety of decision support systems.

Current research and development in the field of biosensors for animal health management focusses on innovative non-invasive, wearable sensors equipped with electronic systems for data collection and transmission of data wirelessly. For specific areas where wearable sensors cannot be multiplexed, especially in disease diagnosis, the development of portable hand-held systems with immediate readout of results will enable fast decision-making processes and help in rapid management of animal diseases. For example, non-invasive screening has been applied to the detection of foot-and-mouth disease using hand-held air samplers with electrostatic particle capture. In this case, infectious viruses are captured and subjected to analysis by real-time PCR (polymerase chain reaction). Such biosensors can hasten the process of monitoring, diagnosis and isolation of contaminated livestock in epidemiological contingencies (Christensen et al., 2011; Wilson, 2015).

On the one hand, in the field, integrated biosensors will enable intensive (frequent and rapid) evaluation of all aspects affecting an animal’s behaviour, genetics and physiology, as well as dynamic changes in metabolism and welfare. On the other hand, they will help manage the production and analysis of the animal’s food composition and help in rapid screening of diseases, which at present cost billions of dollars annually to the livestock industry worldwide. For example, application of nanobiosensors for rapid detection of foot-and-mouth disease in swine has a potential to save costs as well as prevent the spread of infection to the uninfected animals.

Advancements in biosensing technologies is also expected to focus on the use of non-destructive chemical analysis technologies, such as near infrared spectroscopy (NIRS), nuclear magnetic resonance spectroscopy (NMRS) and tunable diode laser absorption spectrometry (TDLAS) for the rapid and detailed evaluation of a biological sample’s chemical composition and safety.

All these exciting technologies need to be developed for seamless performance and validation, before they can be used routinely in research and, ultimately, in a commercial setting. Use of surgically modified animals (e.g., multiple intestinal cannulations, arterials/venous catheters, heart rate telemetry devices) for serial sampling to correlate physiological and metabolic indicators with other variables (e.g., the non-invasive monitoring of behaviour and stress responses, e.g., thermal imaging to assess changes in blood flow). This intensive collection of data from various sensors and complementary analyses will generate a vast magnitude of information, which will need to be effectively collected, compiled, synthesised, securely stored and analysed using a series of advanced statistical, bioinformatics and mathematical modelling approaches. This will require the implementation of a well-integrated and robust data collection, storage and computing infrastructure.

## 2. Monitoring jaw movement of cattle to know the grazing efficiency

Cattle grazing behaviour requires individual monitoring of cattle based on three important parameters, including the location of the animal, analysing animal posture and the movement of the animal, especially movements such as walking and movement of the jaw (Herinaina et al., 2016; Nadin et al., 2012). Jaw movements define the grazing behaviour of the cattle, and there are three different classes of biosensors that can be used to identify such movements. These include:

## 2.1 Mechanical sensors (pressure sensors), acoustic sensors (microphone) and electromyography sensors

Early systems developed to quantify the feeding behaviour of cattle were created exclusively for research purposes. The IGER Behaviour Recorder (Institute of Grassland and Environmental Research, Okehampton, UK) (Rutter et al., 1997) and the ART-MSR pressure sensor (Agroscope Reckenholz-Tänikon ART Research Institute, Modular Signal Recorder MSR145, MSR Electronics GmbH, Switzerland) (Nydegger et al., 2010) were designed specifically to be used in pastures and stables, respectively. The IGER Behaviour Recorder comprises a noseband and an electronic interface connected to a computer to record, analyse and store data at 20Hz (Rutter, 2000). Jaw movement is identified as a pressure peak through the transmission of the movement to the halter and the change in the tube pressure. The installed software can distinguish between bites and chews (Rutter, 2000). Peaks are considered to be bites when they are a combination of a major long peak followed by a smaller sub-peak, or a non-symmetrical peak in the absence of the sub-peak (Nadin et al., 2012). The IGER Behaviour Recorder can also be used to estimate the feed intake with reasonable accuracy, using data on the number of chews and eating duration (using correlation coefficients) (Pahl et al., 2016). Miscalculations in the pressure sensing arise from practical considerations, i.e., due to variations in the tightening of the halter on individual animals, which can result in different pressure values. Another practical issue relates to the measurement of the output wave signal, which can be altered if the halter is mounted too tightly or too loosely. Eating and rumination behaviours in cattle have also been reported with a noseband pressure sensor, consisting of a data logger, incorporated in the noseband to record the jaw movements via a pressure sensor (Braun et al., 2013).

### 2.2 Monitoring jaw movement through acoustic sensing

Acoustic analysis of grazing behaviour has been shown to accurately identify chewing and biting, and therefore can be used to estimate the food intake of cattle (Laca and WallisDeVries, 2000). Acoustic analysis allows differentiation of three types of jaw movements: chew, bite and chew-bite, and microphones can be used to record the jaw sounds of a grazing animal (Ungar and Rutter, 2006). The data can be used to classify ruminating behaviour (Benvenutti et al., 2016; Navon et al., 2013) and are especially helpful in monitoring animal wellbeing.

Acoustic sensing of jaw movements can be classified based on the microphone location and acoustic system classification. Detailed analysis of the systems currently in use are elaborated in Figure 2. Some systems are simple and detect jaw movements based on 10-minute recordings of grazing sessions on a camera, with an accuracy of 94% (Herinaina et al., 2016; Navon et al.,2013). Signal patterns are analysed by the machine-learning algorithms to determine intervals between jaw movements, intensity of each jaw movement (observed as a peak in the time domain), their duration, and their integration in a sequence of behaviours (Navon et al., 2013). The discrimination is based on the signal patterns produced during biting and chewing in a 1- kHz sound window: peak frequency, peak intensity, average intensity and their duration (Laca et al., 2000). Clapham and colleagues have demonstrated a fully automatic Chew-Bite Real-Time Algorithm for detection and classification of ingestive events during cattle grazing. The system consists of a directional wide-frequency microphone facing inwards on the forehead of the animal, and coupled with the signal analysis and decision logic algorithm, it can detect and analyse bites and chews with an accuracy of 94% (Clapham et al., 2011). Milone *et al*. have reported the detection and classification of bites, chews and chew-bites with the help of the Hidden Markov model, which estimates the sequences of bites, chews and chew-bites using acoustic spectrum characteristics like decibels, by each sound. The successful classification is reported to range between 61% and 99% (Milone et al., 2012).

**Figure 2.**
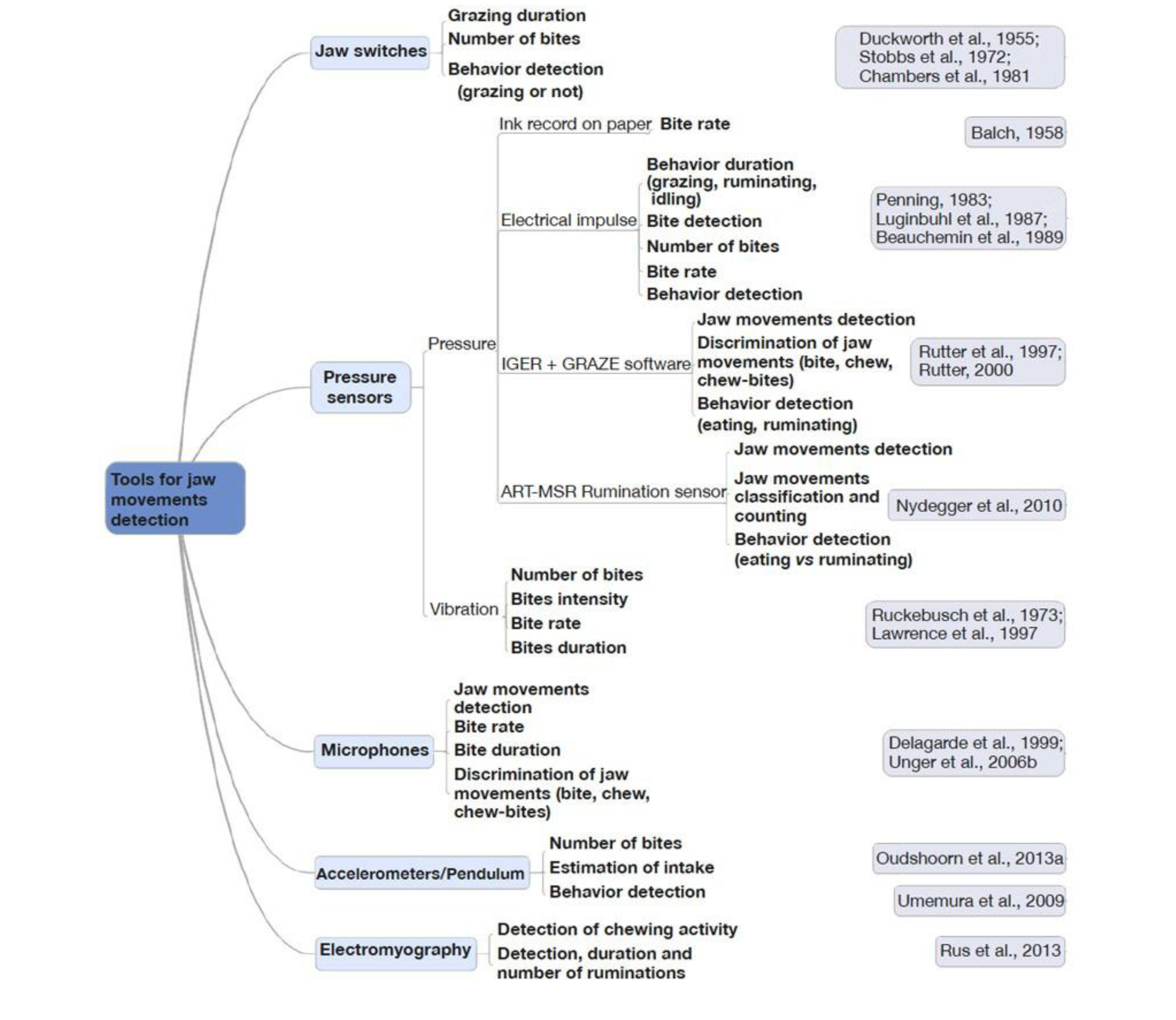
Primary tools used to detect jaw movements in cattle. Source: (Herinaina et al., 2016)

Microphone-based methods have demonstrated good accuracy for the detection of jaw movements and can differentiate between three different kinds of jaw movements. However, one of the key disadvantages of acoustic systems in outdoor environments is their susceptibility to environmental noises. Acoustic interpretation techniques that can overcome these disturbances still need to be advanced before these systems are deployed on the farm.

### 2.3 Acceleration sensors for jaw movement and feeding behaviour

Accelerometer sensors convert physical acceleration recorded from motion or gravity into a voltage output. Accelerometers can be used to measure static acceleration due to gravity, the low-frequency component of the acceleration and the dynamic acceleration due to animal movement (Herinaina et al., 2016). Several researchers have demonstrated the use of accelerometers for analysing the grazing behaviour of animals (Mattachini et al., 2016; Tani et al., 2013; Giovanetti et al., 2017). Andriamandroso *et al*. (Andriamandroso et al., 2015) used smartphone inertial measurement units (IMUs) to count the number of bites through a frequency pattern of single-axis acceleration data.

Oudshoorn *et al*. (Oudshoorn et al., 2013) were the first to use a 3-axis accelerometer to quantify cow bites. The method involved visualisation of the recorded signals from the three individual orthogonal axes to determine the signal that best matched the recorded bites. The method, however, indicated an average correlation coefficient of 0.65. Umemura *et al*. were able to monitor the jaw movements by modifying a pedometer into a pendulum, attached to the lower jaw. The data from the device could be downloaded wirelessly and the system showed 90% accuracy in measuring jaw movements when compared to manual counts over 10-minute segments (Umemura et al., 2009).

One key issue associated with the use of accelerometers is the sensitivity of the signals recorded. Undesirable signals due to rapid head movement or the movement of ears can interfere with data interpretation, and the system would require a pre-processing of the signal relative to the jaw movements in order to be useful for precision livestock farming. Interestingly, the IMUs, which combine several sensors such as the accelerometers, gyroscope, magnetometer and GPS, can offer a real advantage in terms of multiple-parameter measurements related to animal feeding behaviour and animal position (active or not). Figure 3 details the key components of multi-parametric sensors deployed for precision livestock farming. These multi-parameter sensors collect and integrate the data from individual sensors to provide a comprehensive health picture of individual animals, as well as herd behaviour. Such integrated sensors will also enable predictions on animal health events and disease. In the event of a disease outbreak, multi-parameter sensors can assist in identification and isolation of affected livestock before the spread of the outbreak and potentially prevent unnecessary culling of uninfected animals, as is the current practice in animal husbandry.

**Figure 3.**
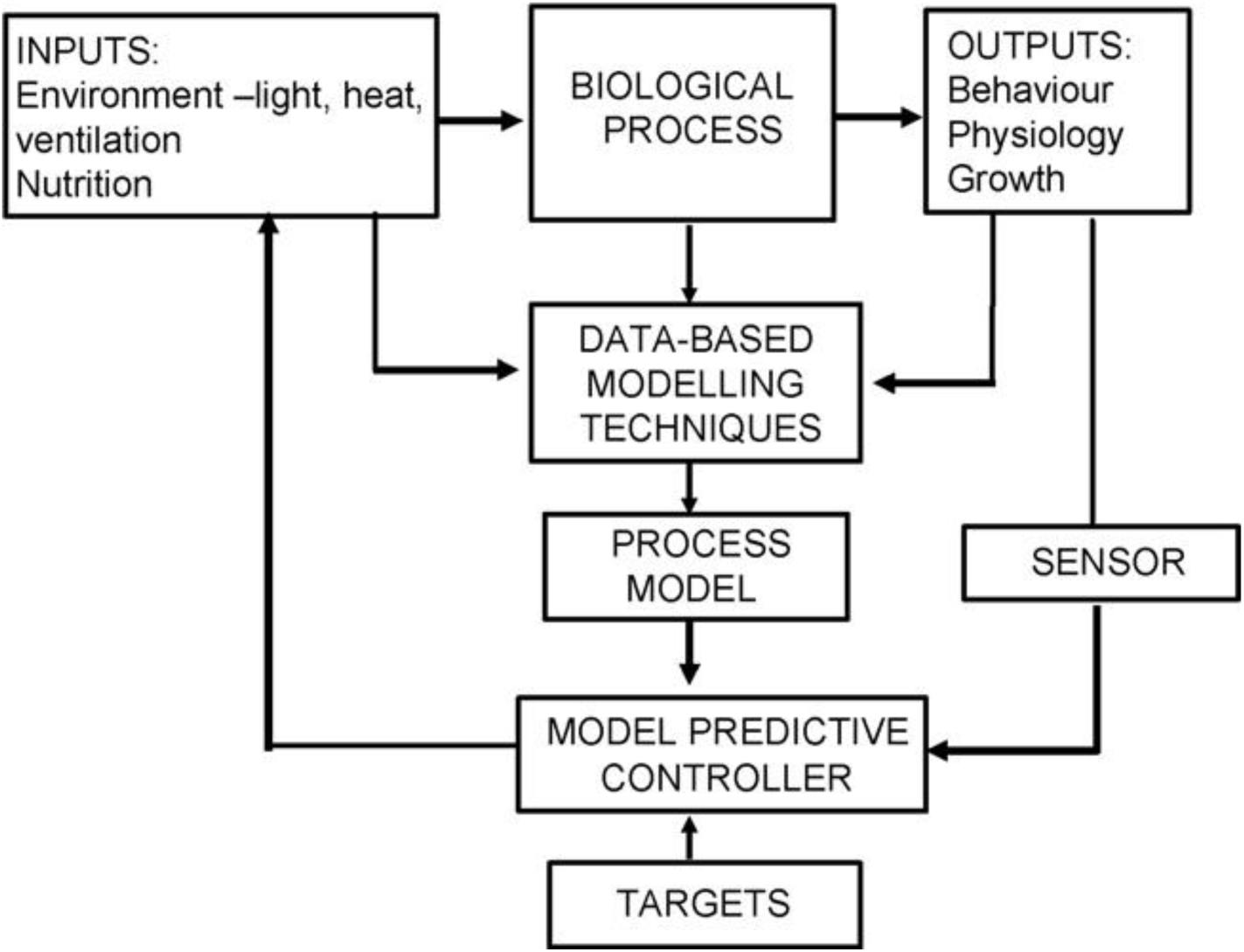
Schematic overview of the key components of Precision Livestock Farming to control biological processes. Source: (Wathes et al., 2008) Reprinted with permission from Elsevier Ltd.

## 3. Biosensors for breath analysis

Disease diagnosis by identification of volatile organic compounds (VOCs) has long been of interest to researchers, as it offers a non-invasive methodology. VOCs can be found in the breath, blood, faeces, skin, urine and vaginal fluids of animals as well as humans (Burciaga-Robles et al., 2009; Garner et al., 2009; Spinhirne et al., 2004). These compounds are produced by a number of biochemical reactions, pathogens, and host pathogen interactions and are affected by a number of biological variables such as age, actions, and biochemical pathways (Sethi et al., 2013).Breath monitoring provides a non-invasive and easy approach to determine the physiological and general health status of animals. Advances in sampling methods, like solid-phase and needle trap micro-extraction, and developments in techniques for representative breath sampling (Turner et al., 2012) can be coupled with modern analytical technologies (spectroscopy techniques and electronic noses) to allow for precise analysis of breath composition at an unprecedented level (Pereira et al., 2015b). One of the key challenges that require important attention is the statistical analysis and data interpretation of large and potentially heterogeneous datasets collected from research on the exhaled breath composition from animals.

Metabolites in the breath include gasses like hydrogen and methane and volatile organic compounds such as fatty acids, which can act as biomarkers for metabolic and pathologic processes. Usually, the glucose level in blood is associated with VOCs like ketone bodies, ethanol, methanol and exogenous compounds (Leopold et al., 2014).

In cattle, analysis of VOCs has been explored to diagnose bovine respiratory disease (Burciaga-Robles et al., 2009), brucellosis (Knobloch et al., 2009), bovine tuberculosis (Fend et al., 2005; Peled et al., 2012), Johne's disease (Kumanan et al., 2009), ketoacidosis (Mottram et al., 1999), and normal rumen physiology. A rapid, non-invasive identification of foot-and-mouth disease has been performed using air samples collected with a hand-held prototype device equipped with electrostatic particle capture in a microchip chamber of 10-15 μL (Christensen et al., 2011).

## 4. Sensors analysing metabolites in perspiration

Most biosensors developed for analysing metabolites in sweat were developed with the purpose of human health monitoring. These have been used to analyse sodium concentration (Schazmann et al., 2010) and lactate levels, and converted to portable formats (belt form) to analyse sweat. The electrochemical sensor for lactate levels includes a flexible printed tattoo that can detect lactate levels with linearity up to 20 mM. The sensor has been shown to be resilient against mechanical deformation. This sensor can also be adapted for use in animal sweat monitoring, especially as a sign of physical stress in animals (Jia et al., 2013). Others have developed an adhesive radio-frequency identification (RFID) sensor patch, which allows for potentiometric sensing of solutes and surface temperature that can be read on a smartphone application (Rose et al., 2015). New research in this area has led to the development of a fully integrated wearable sensor array for multiplexed analysis of perspiration. The system includes a mechanically flexible sensor array that measures metabolites like glucose and lactate, and electrolyte composition such as sodium and potassium ions. The sensor also integrates temperature measurement for comprehensive analysis (Gao et al., 2016).

Bovine tuberculosis (*M bovis*) is a chronic bacterial disease affecting cattle and can occasionally spread to humans by the inhalation of aerosols or consumption of unpasteurised milk. The ability to identify volatile organic compounds produced by pathogens has been applied to this technology for the detection of *M*. *bovis* infection by analysing the changes in the volatile organic compound profiles present in breath. More recently, a proof of concept has been presented to reveal that the breath-derived volatile organic compound analysis can be used to differentiate between healthy and *M. bovis*-infected cattle (Ellis et al., 2014).

The TB Breathalyser system for tuberculosis diagnosis in humans, developed by Rapid Biosensor Systems, is already available in the market (McNerney et al., 2010). The gas-chromatography/mass-spectrometry analysis has revealed the presence of VOCs associated with *M. bovis* infection. A nanotechnology-based array of sensors has been tailored for detection of *M. bovis*-infected cattle via breath, which allows real-time cattle monitoring (Peled et al., 2012). Kumanan *et al*. have reported the development of a membrane-strip-based lateral-flow biosensor combined with a high-throughput microtiter plate assay to enable highly sensitive reverse transcriptase polymerase chain reaction (RT-PCR) -based detection of viable *Mycobacterium* (*M*.) *avium* subsp. paratuberculosis cells in faecal samples (Kumanan et al., 2009).

## 5. Analysis of tears for continuous glucose monitoring

Metabolites in tears can provide information about the concentration of these metabolites in blood and provide a non-invasive continuous monitoring technique. Iguchi *et al*. have reported the development of a flexible, wearable amperometric glucose sensor using immobilised glucose oxidase on a flexible oxygen electrode (Pt working electrode and Ag/AgCl counter/reference electrode). The biosensor is fabricated using Soft-MEMS techniques onto a functional polymer membrane (Iguchi et al., 2007). Others are working towards the development of a biosensor for self-monitoring of tear glucose and are currently in the animal testing stages (La Belle et al.,2014) (Yonemori et al., 2009).

## 6. *In vivo* implanted biosensor to analyse stress in fish

Fish health is affected by multiple environmental parameters as well as conditions in the fish farms. Stressors include water pollution and changes in climate. Farm management practices like stocking density and water exchange can also induce fish stress (Figure 4). Wu *et al*. have devised an implantable biosensor that detects the composition of eyeball scleral interstitial fluid in fish. The contents of the fluid correlate well with their concentrations in blood. Stress due to changes in water chemistry, dissolved oxygen content, pH, and metal toxicity were monitored, and behavioural changes such as attacking behaviour and visual irritation were recorded (Wu et al., 2015). Hibi *et al*. have developed a wireless biosensor system for continuous monitoring of stress biomarker L-lactic acid in fish using the eyeball interstitial sclera fluid site for sensor implantation. The biosensor allows for wireless monitoring of L-lactic acid in free-swimming fish (Hibi et al., 2012).

**Figure 4.**
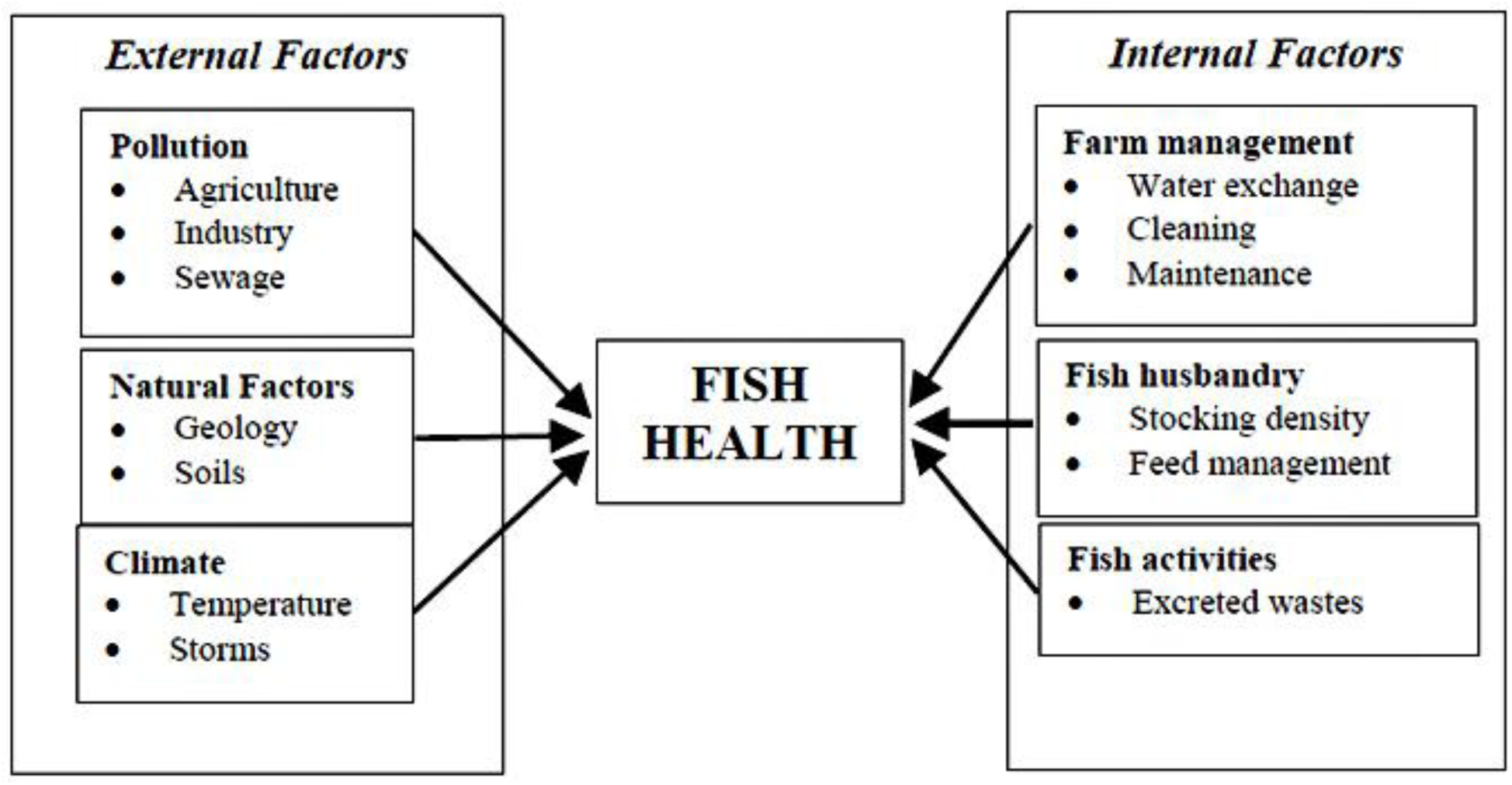
Environmental and farm management practices affecting fish health. Source: (Ingram et al., 2005)

## 7. Detection of ovulation

### 7.1 Progesterone

Breeding forms an integral part of livestock farming. Detection of the ovulation period in cattle is important in order to determine the time window for artificial insemination. Conventional oestrus detection involves ocular inspection of cattle by skilled labour, which is expensive as well as inefficient. Biosensors for ovulation detection have been researched for a long time. Pemberton and colleagues reported a device able to determine ovulation using a disposable screen-printed amperometric progesterone biosensor, operated in a competitive immunoassay. The biosensor included a monoclonal anti-progesterone antibody (mAb) immobilised on a screen-printed carbon electrode (SPCE). It was later incorporated into a thin-layer flow cell offering advantages such as on-line analysis and improved fluid handling with the possibility of future automation (Pemberton et al., 2001). The Herd Navigation^®^ system was developed in 2008 for commercial use and combines five sensing systems, including progesterone in the milk. Herd Navigator™ (Durkin and DeLaval, 2010) measures the level of progesterone in milk and the software suggests the insemination time, lists animals for final pregnancy confirmation, indicates early abortion and lists the cows at risk for cysts and prolonged anoestrus (Mazeris, 2010). Oestrus detection rates of 95-97% have been reported in the farms in Denmark, with significantly higher pregnancy rates (up to 42-50%) than the conventional techniques (Blom and Ridder, 2010; Vreeburg, 2010b). There are also reports indicating cost savings of €250 and €350/cow per year, as farmers do not have to spend money on expensive pregnancy tests (Mazeris, 2010; Leonardi et al., 2013).

### 7.2 SPR-based biosensors for progesterone

More recently, Zeidan and colleagues have reported the development of a progesterone sensor by integrating novel aptamer development with a nanoEnhanced Surface Plasmon Resonance imaging sensor (SPRi). The authors first developed X-aptamers and selected them for binding to progesterone. Then, the multi-array feature of SPRi was used to develop an optimised biosensor capable of simultaneously screening the 9 X-aptamers for binding affinities. The sensor surface was further optimised in a sandwich assay, where nanoEnhancers (NIR-streptavidin-coated quantum dots) were used for ultrasensitive detection of progesterone molecules (Zeidan et al., 2016).

### 7.3 Herd Navigator ™ for monitoring ovulation

Herd Navigator has excellent oestrus detection rates, but the system is too expensive. Aiming at improving detection reliability using low-cost sensor data, Jónsson *et al*. reported the combination of information from step count and leg tilt sensors. The authors developed a change-detection algorithm that can analyse cow-specific data in real time. The system has shown an increase in the successful alerts and significantly reduced false positives (Jónsson et al., 2011).

### 7.4 Intravaginal probes

Andersson *et al*. tested a wireless intravaginal probe, with a possibility for automation of the process. The probe is based on the measurements of conductivity and temperature, and also senses the movement of the animal. These parameters can all be used independently to detect oestrus. Although still in the testing phase, the device has been shown to have a higher reliability and to be more resistant to external disturbances as compared to existing alternatives (Andersson et al., 2016; Andersson et al., 2015).

## 8 Biosensors for animal diseases

### 8.1 Bovine Respiratory Disease

Bovine Herpes Virus-1 (BHV-1) is a major viral pathogen of Bovine Respiratory Disease (BRD), the prominent cause of economic loss ($2 billion annually in the US alone) to the cattle and dairy industries. Tarasov *et al*. report the development of an extended-gate field-effect transistor (FET) for direct potentiometric serological diagnosis of the BHV-1 viral protein via an IgE-coated immunosensor. The biosensor was presented to be sensitive and selective to anti-IgE present in commercially available anti-BHV-1 antiserum and in real serum samples from cattle. The system was shown to be faster than the traditionally used ELISA, amenable to multiplexing, and easily integrated into POC devices (Tarasov et al., 2016).

Schaefer and colleagues have investigated the use of an automated, RFID-driven, infrared thermography technology to determine BRD in cattle. The animals were monitored for BRD using biometric clinical scores, body temperature, haematology, serum cortisol and infrared thermal values. The data collected showed a correlation between animals positive for BRD with higher peak infrared thermal values of 35.7 ± 0.35 °C, in comparison to the true negative animals’ 34.9 ± 0.22 °C. The study is a proof of concept that the thermography data could be non-invasively and automatically collected on the basis of a system developed around the animals’ water station (Schaefer et al., 2012).

### 8.2 Detection of Bovine viral diarrhoea virus (BVDV)

While ELISA and PCR-based methods have long been used for the detection of BVDV (Da Silva et al., 1995; Pritchard et al., 2002), rapid detection of BVDV requires an on-site monitoring and detection system to expedite the diagnosis and minimise the spread of disease in the herd. BVD disease affects beef and dairy industries worldwide, with severe implications to costs. Rapid diagnosis of BVDV through on-farm analysis is critical for herd protection and prevention of herd outbreaks. To this end, Montrose *et al*. have developed a fully integrated nanowire-based immunosensor to detect BVDV in serum. The biosensor has BVD virus as a capture molecule, which is covalently immobilised to a polymer electrodeposited onto a nanowire (Montrose et al. 2015). Luo *et al*. have developed an electrospun biosensor based on capillary separation and conductometric immunoassay for the detection of BVDV antibodies. The detection time of the biosensor is 8 minutes, and the detection limit is 10^3^ CCID/mL for BVDV viral samples (Luo et al., 2010). Heinze *et al*. have utilised microparticle immunoagglutination assays on a microfluidic chip using forward light scattering measurements to detect BVDV particles (Heinze et al., 2009).

### 8.3 Avian influenza virus

Avian Influenza Virus (AIV) infections have been a major cause of mortality, and rapid detection methods for avian influenza have huge clinical, economical and epidemiological implications. Diagnosis with ELISA and PCR is generally time-consuming and expensive, requiring transport of samples to specialised laboratories. In recent years, there have been several developments to miniaturise as well as provide assays that can be used on the farms for diagnosis of disease. Diouani *et al*. and Wang *et al*. have reported the development of a miniaturised gold electrode biosensor using impedance spectroscopy to detect H_7_N_1_. The biosensor is based on the detection of immobilised H_7_N_1_ antibodies onto a bio-functionalised gold electrode (Diouani et al., 2008) (Wang et al., 2009).

Xu *et al*. have developed an interferometric biosensor immunoassay for the direct and label-free detection of avian influenza strains H_7_ (two strains) and H_8_ (one strain) through whole virus capture on a planar optical waveguide. The assay relies on the index of refractive changes occurring upon binding of virus particles to unique antigen-specific (hemagglutinin) antibodies on the waveguide surface (Xu et al., 2007). Others have developed DNA-aptamers as recognition elements in portable Surface Plasmon Resonance (SPR) -based biosensors for rapid detection of H_5_N_1_ in swab samples from poultry (Bai et al., 2012). Luminescence resonance energy transfer (LRET) -based biosensors for the ultrasensitive detection of the H_7_ strain (Ye et al., 2014) and indium-tin-oxide thin-film transistors (ITO TFTs) on a glass substrate for immune detection of H_5_N_1_ antibodies have also been reported (Guo et al., 2013).

Others have developed quartz crystal microbalance (QCM) aptasensors based on ssDNA crosslinked polymeric hydrogel for rapid, sensitive and specific detection of H_5_N_1_ within 30 minutes (Wang and Li, 2013) and QCM-based immunosensors to detect H_5_N_1_ (Li et al., 2011). Impedance-based sensitive and rapid methods for screening for the H_5_ subtype using immune-magnetic nanoparticles have been reported, in which the virus is separated and the measurement of an interdigitated microelectrode utilised for impedance measurement (Lum et al., 2012).

### 8.4 Foot-and-mouth disease

Rapid initial diagnosis of foot-and-mouth disease virus (FMDV) is essential for faster diagnosis. Several biosensors have been developed recently to provide portable systems for the diagnosis of FMDV. These have been reviewed extensively by Niedbalski (Niedbalski, 2016). The systems developed include lateral flow immunochromatographic (LFI) for the detection of antibodies against FMDV proteins (Yang et al., 2015; Yang et al., 2013) to detect FMDV serotypes O, A, Asia 1, SAT 2 and non-serotype-specific FMDV. Several FMD-specific real-time RT-PCR (rRT-PCR) assays have been made into portable mobile platforms for in-field detection of FDMV. These include the Cepheid Smart Cycler Real-time PCR machine (Hearps et al., 2002), and the BioSeeq-Vet (Smiths Detection). Genie I, a portable platform, also allows for the on-site detection of viral RNA by reverse-transcription loop-mediated isothermal amplification (RT-LAMP) (Waters et al., 2014). Recently, rapid identification of FMDV has been reported using SpectroSens™ optical microchip sensors. Selective identification of FMDV is conducted in minutes and displayed as a yes/no readout using a hand-held device (Bhatta et al., 2012). Infrared thermography (IRT), a quantitative method for the assessment of body surface temperature, can be useful for the early detection of FMDV in the field. Temperature screening can be used to isolate potentially sick animals at an early stage and prevent the spread of disease. Microarrays, designed for laboratory diagnosis of FMD, offer greater screening capabilities for FMDV detection and can be regarded as an alternative to classical diagnostic methods. However, the apparatus needs to be miniaturised and made portable before it can be used directly in the field.

### 8.5 Automated Detection of Mastitis

Mastitis is associated with the inflammation of the udder in cattle due to an infection by *Staphylococcus aureus*. Mastitis detection in milk is based on two milk quality aspects: the somatic cell count (SCC) and the presence of visibly abnormal milk in the case of clinical mastitis. Efficient detection of mastitis is essential in order to manage the infected cattle and progression to clinical mastitis (Hogeveen et al., 2010). Neitzel and colleagues have developed an indirect on-line sensor system based on the automated California Mastitis Test (CMT) in milk (Neitzel et al., 2014). Duarte *et al*. have reported the development of an immune assay based on coupling with magnetic nanoparticles, which is analysed using a lab-on-a-chip magneto resistive cytometer, with microfluidic sample handling (Duarte et al., 2016). Others have reported the development of selective amperometric biosensors for infected milk detection based on the quantification of the catalase enzyme, which is immobilised on a thin-layer enzyme cell (Fűtő et al., 2012).

### 8.6 Subclinical ketosis

Nanobiosensors can significantly aid in the real-time detection of beta-hydroxy butyrate from blood or milk to assessthe energy balance of the animals¨ β-hydroxybutyrate (βΗΒΑ) is an indicator of subclinical ketosis, a common disease in dairy cows. Subclinical ketosis is one of the metabolic diseases associated with negative energy balance during the transition period, as well as decreased milk yields, impaired reproductive performance and higher risk of clinical ketosis, resulting in economic losses (Ospina et al., 2010). Weng and colleagues have recently reported the development of the on-chip detection of βΗΒΑ using a miniaturised, cost-effective optical sensor. The authors report that the analysis can be completed in 1 minute and has a detection limit of 0.05 mM βΗΒΑ (Weng et al., 2015b). In another study, a biosensor using quantum dots (QDs) modified with cofactor nicotinamide adenine dinucleotide (NAD+) has been used for sensing βΗΒΑ concentration in a cow's blood and milk sample. The detection is performed on a custom-designed microfluidic platform combined with a low-cost, miniaturised optical sensor. The sensing platform has a detection limit better than the previous method at 35 μM (Weng et al., 2015a) (Neethirajan et al., 2016).

### 8.7. Detection of porcine reproductive and respiratory syndrome (PRRS) virus

Infection with the porcine reproductive and respiratory syndrome virus (PRRSV) results in PRRS in pigs, also known as the blue-ear pig disease, causing reproductive failure in breeding stock accompanied by respiratory tract illness in piglets. The disease costs the United States swine industry around $644 million annually according to a 2011 study (Holtkamp et al., 2013), and recent estimates in Europe found that it cost almost 1.5b€ in the year 2013. Several immunodetection-based biosensors have been reported for the detection of PRRS and are detailed in Table 1.

**Table 1.**
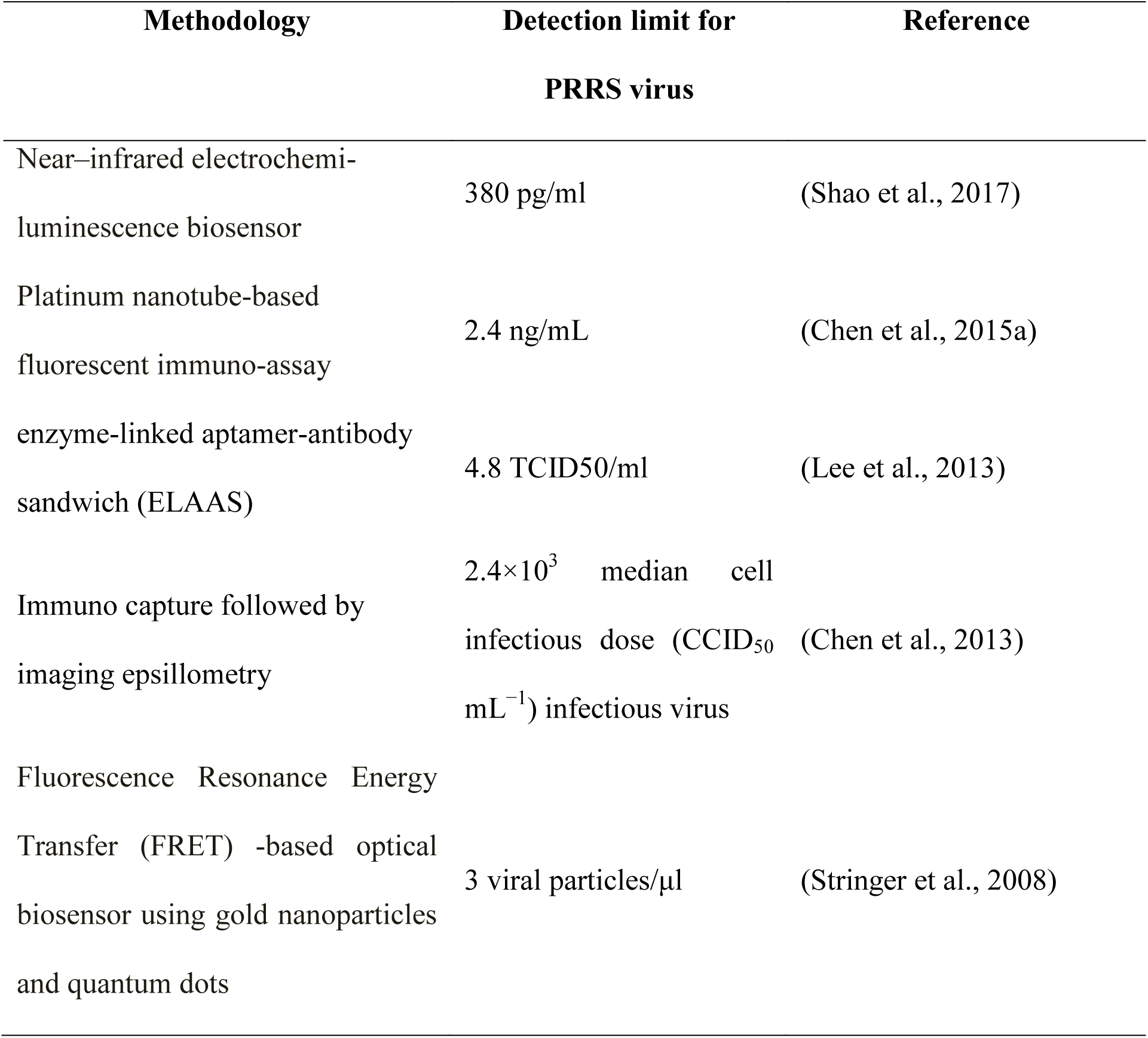
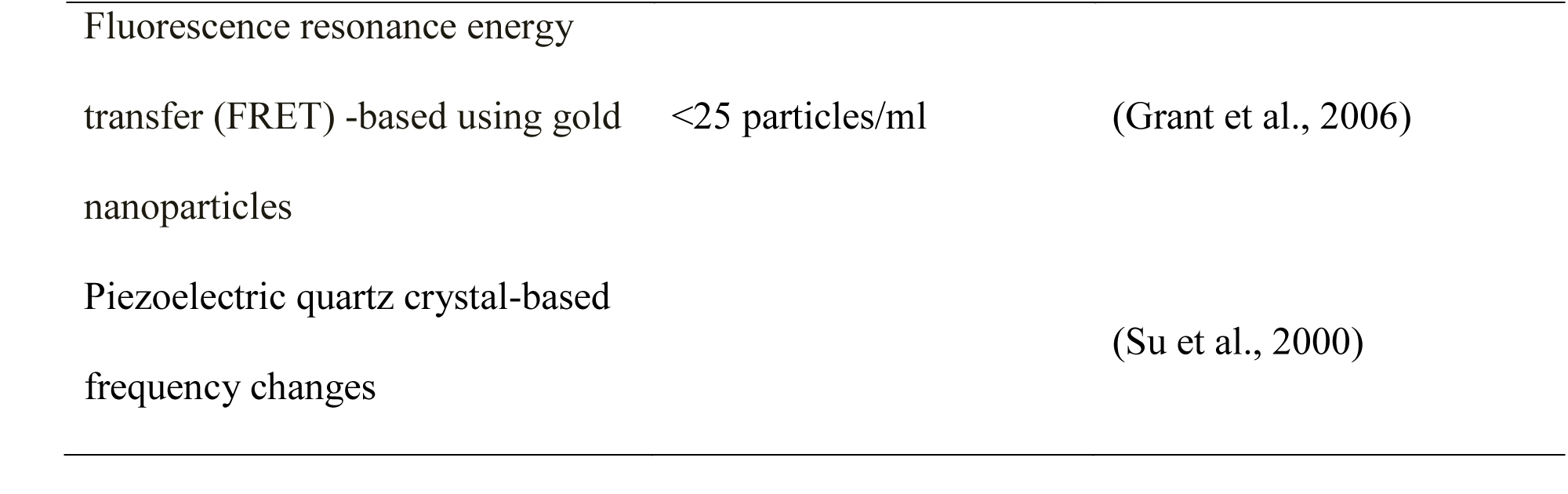
*Techniques used for the detection of porcine reproductive and respiratory syndrome virus*

**Table 2.**
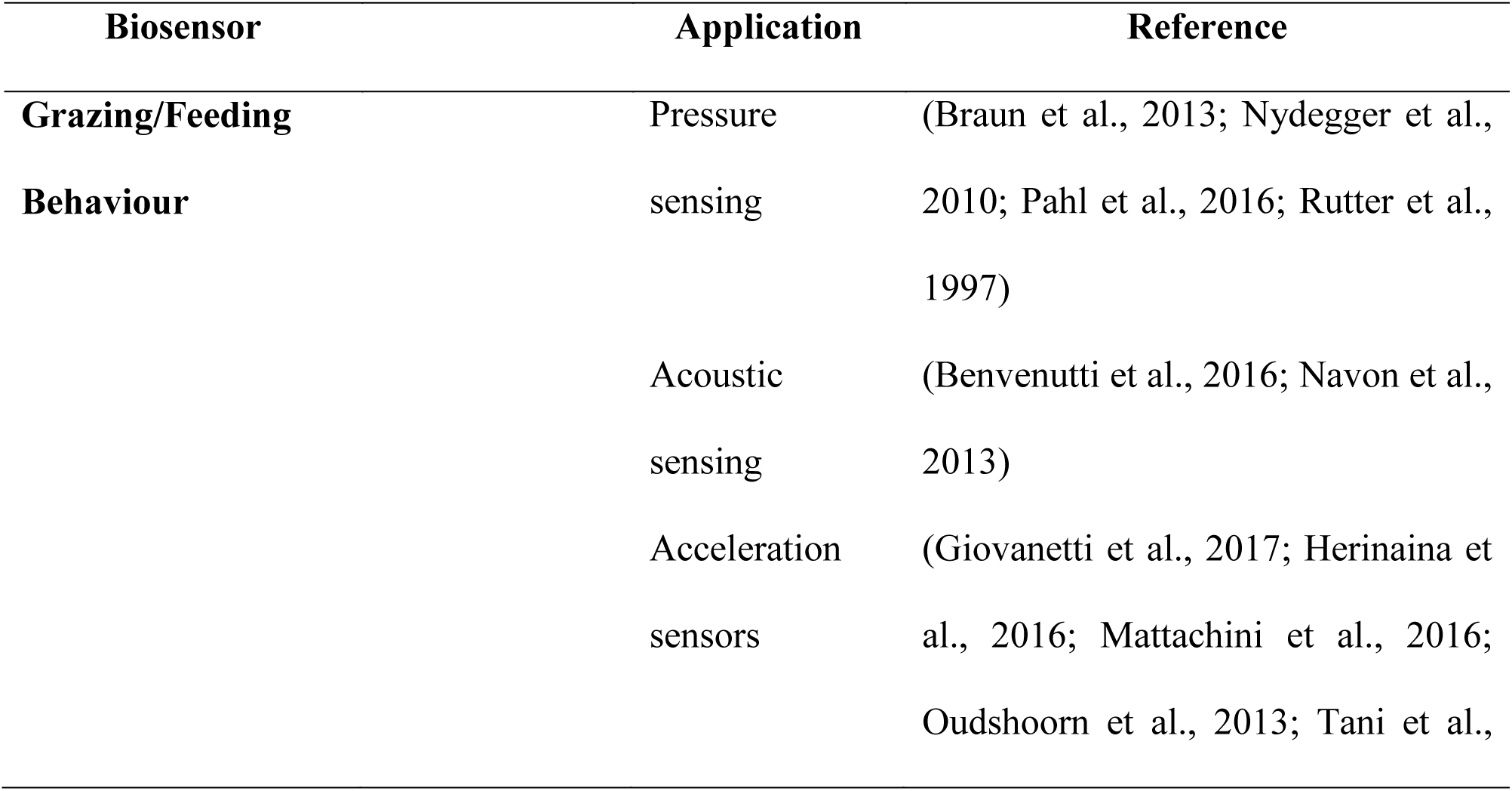
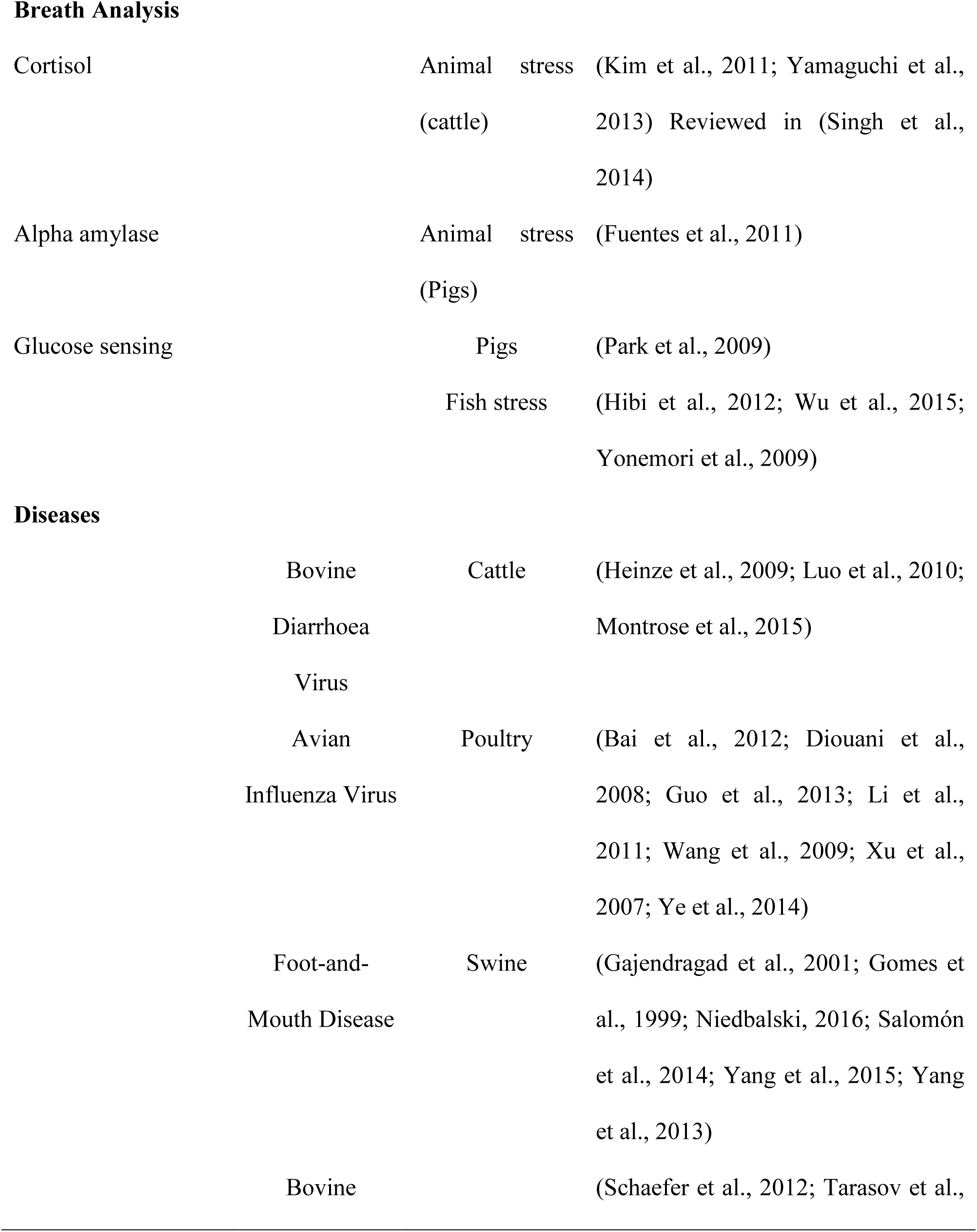
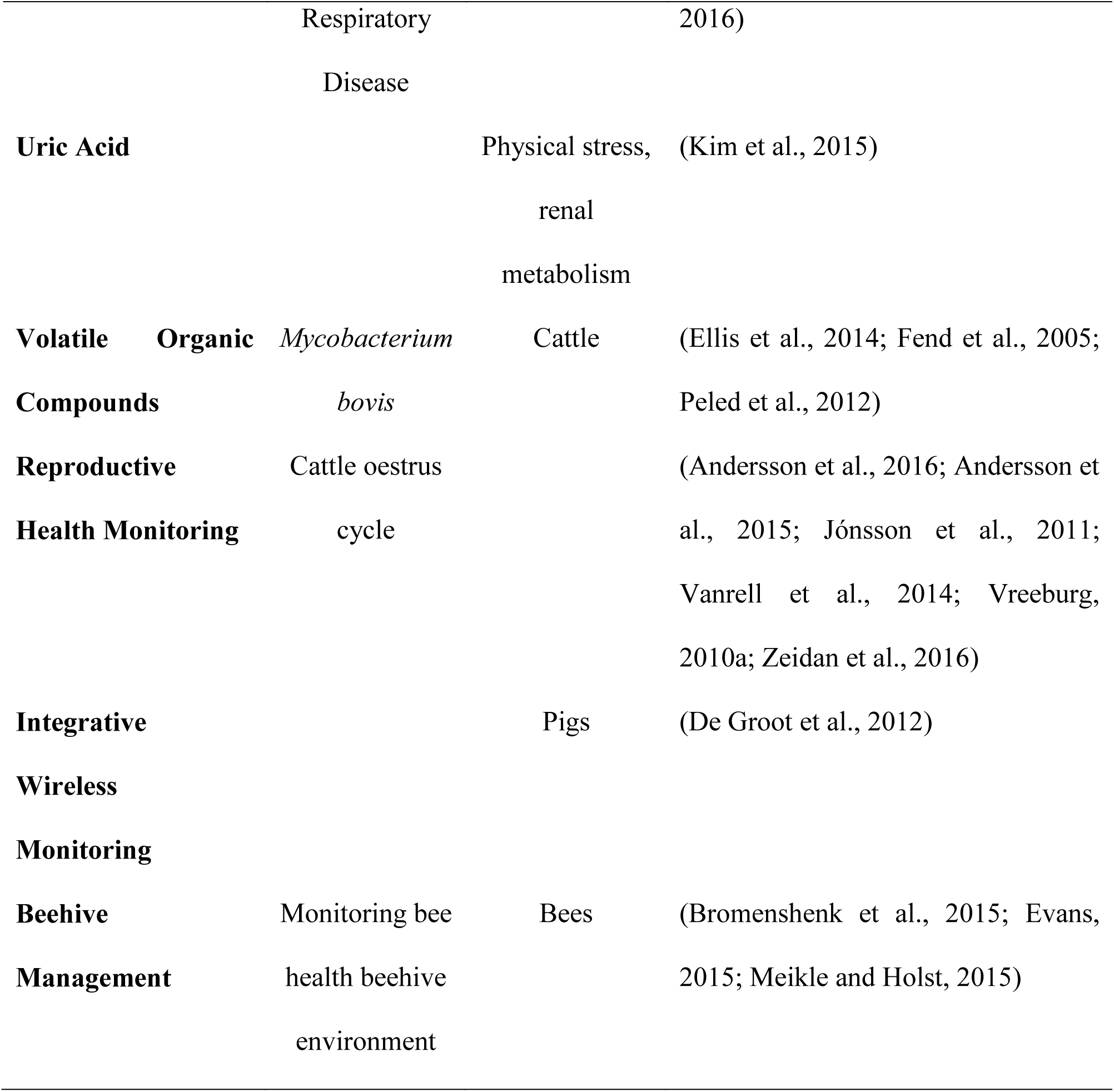
*Biosensor applications in animal health monitoring*

**Table 3.**
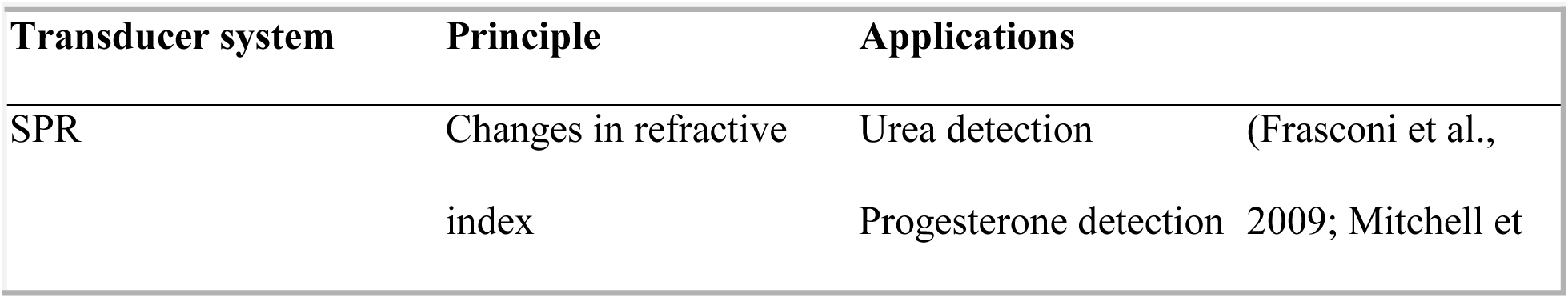
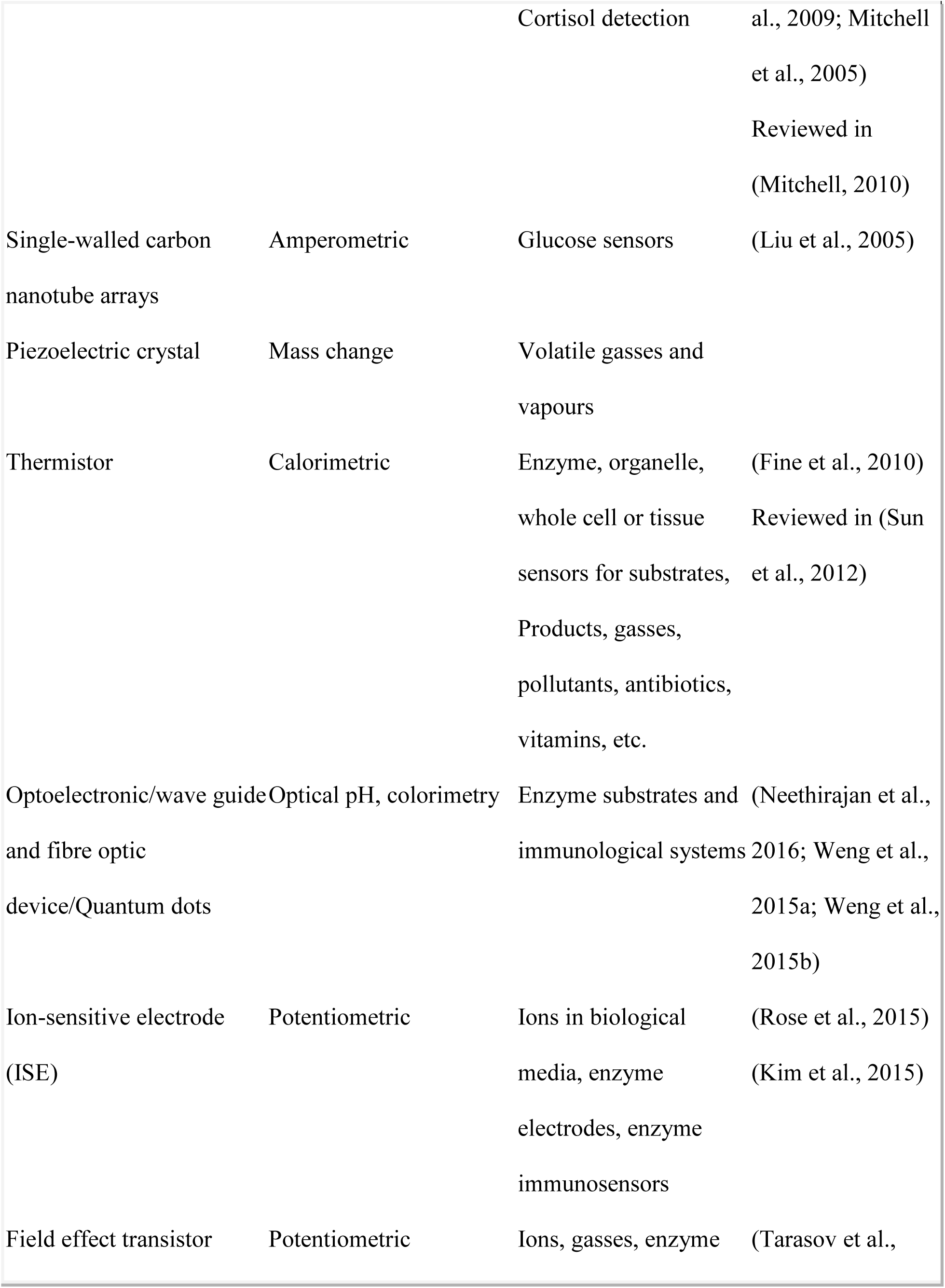
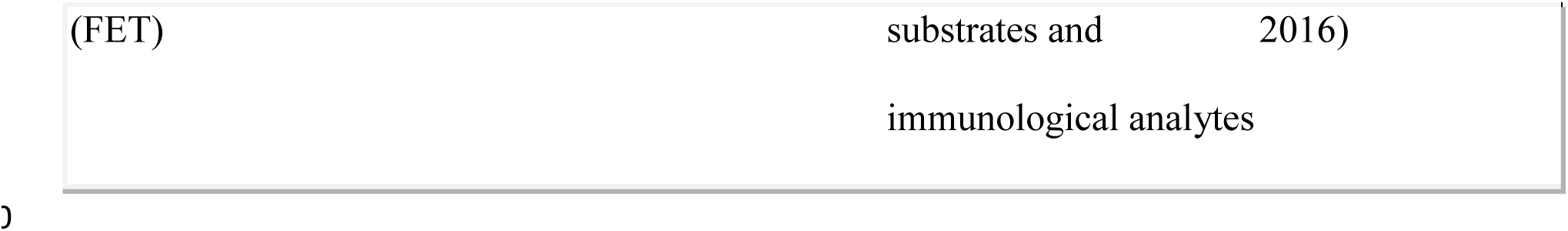
*Biosensor Transducers and Applications*

### 8.8. Salivary detection of metabolites of clinical significance

Saliva sampling for disease and other biochemical markers of physiological health is an attractive alternative to blood sampling, as it is non-invasive in nature (Bandodkar and Wang, 2014).The method is particularly useful for animal monitoring and disease diagnostics, as blood collection from animals is considered to be a stress inducer and may have an impact on the biochemical parameters being diagnosed. The ability to collect and immediately analyse the salivary samples on-site provides numerous advantages for field applications. Biomarkers in saliva can be helpful in numerous ways, e.g.: (i) early detection and diagnosis of diseases; (ii) in supporting the decision-making processes for animal handling; and (iii) to monitor the progression of disease (Malon et al., 2014). However, it must be noted that current analysis procedures, if applied to saliva, would require huge amounts of salivary probes for the biochemical assays. Although saliva sampling using oral fluid collectors and commercial devices (Mottram et al., 2004) is generally safe and convenient to use and provides a sufficient homogeneous sample with low viscosity, it still presents several shortcomings, such as (i) the requirement of supervision; (ii) the need to follow the procedures carefully to ensure sample adequacy; and (iii) it is a time-consuming process. Moreover, the assays for biomarkers in saliva have to be calibrated against the assays for blood samples to ensure the sensitivity of detection and the robustness of the assays.

#### 8.8.1 Biosensors for salivary uric acid

An abnormal concentration of uric acid acts as a biomarker for several diseases, such as metabolic syndrome, renal syndrome, and abnormalities in purine metabolism (Nakagawa et al., 2006; Nyhan, 1997). Uric acid is also known to be present in response to physical stress (Hellsten et al., 1997). Detection of uric acid in saliva presents a non-invasive method, and there is a good correlation between uric acid levels in blood and saliva (Soukup et al., 2012). Kim *et al*. have reported the development of a wearable salivary uric acid mouth guard sensor in which a uricase-modified screen-printed electrode system is integrated into a mouth guard platform. It uses miniaturised instrumentation electronics featuring a potentiostat, a microcontroller, and a Bluetooth Low Energy (BLE) transceiver. This platform enables real-time wireless transmission of information to standard smartphones and other storage devices (Kim et al., 2015).

#### 8.8.2 Measurement of salivary cortisol as an animal stress biomarker

Measurement of corticosteroid hormones is commonly used as a biomarker of an animal's response to stress. The difficulties in obtaining blood samples and the recognition of the stressor effect of blood sampling are primary drivers for the use of minimally invasive sample media. Salivary cortisol has been established as a viable indicator of stress levels in animals held in captive environments. The suitability of a cortisol assay and its validation have been detailed in the review by Cook (Cook, 2012). More recently, Yamaguchi *et al*. have demonstrated the development of a cortisol immunosensor for the non-invasive and quantitative analysis of salivary cortisol. The immunosensor detects current resulting from a competitive reaction between the sample cortisol and a glucose oxidase (GOD)-labelled cortisol conjugate, and quantifies cortisol levels based on a calibration curve. The technique takes 35 minutes for analysing salivary cortisol levels and the device can be used on-site. The method was also shown to closely correspond to the currently available ELISA method (Yamaguchi et al., 2013).

#### 8.8.3 Measurement of salivary glucose

Salivary glucose biosensors have primarily been developed for human use and for domestic pets (Reusch et al., 2006; Stein and Greco, 2002), but they have vast potential in monitoring livestock health. Most of the sensors developed for humans have also been tested in animals. The sensors are lightweight and portable and can be used directly on-field (Park et al., 2009).

### 8.9. In vivo real-time sensing for uric acid in poultry

Energy pathways in birds are lipid metabolism rather than the glucose metabolism dominant in other animals, and uric acid levels are indicative of protein catabolism in birds (Gumus et al., 2014). Numerous methods for uric acid detection based on chemiluminescence, spectrophotometry, fluorescence (Martinez-Pérez et al., 2003) and electrochemistry (Jindal et al., 2012). have been developed. However, Gumus *et al*. have reported the development of uric acid sensing based on uricase enzyme for *in vivo* applications. It is an enzyme-based method using Pt/Ir wire and Ag/AgCl paste. The sensor has a linear response for uric acid in the range of 0.05 to 0.6 mM uric acid, which covers the physiological levels for avian species and is able to transmit the data wirelessly.

### 8.10. Salivary alpha amylase as a stress biomarker in pigs

While cortisol is an essential hormone responsible for the regulation of stress, salivary alpha-amylase is a novel biomarker for psycho-social stress. This has been validated by Fuentes *et al*. using an automated spectrophotometric method for salivary alpha-amylase measurement (Fuentes et al., 2011). Wu *et al*. have fabricated an inexpensive, disposable α-amylase biosensor by immobilising a layer of starch gel on a thick-film magnetoelastic sensor. When exposed to α- amylase, the resonance frequency of the starch gel-immobilised sensor increases in proportion to the starch hydrolysis by α-amylase, allowing for its quantification. The sensor described can detect 75 to 125 U/ml α-amylase (Wu et al., 2007).

### 9. Livestock monitoring systems for observing physiological parameters and health of cattle

Jegadeesan *et al*. have proposed a two-component system. The first component is the monitoring and collection of data on the health parameters of animals in the field, and the second component is the monitoring and acquisition of data on animals from the farms. Animals are subjected to a variety of stress factors during their lives on farms. These include stressors due to changes in temperature, transport across farms, physiological stress due to ill health or improper food intake as well as stress due to restraint (detailed in Figure 5). This information on the external environment and animal health can be collected and analysed in real time using closed-circuit cameras. The complete system is expected to work independently, making necessary changes in response to the real-time data inputs. Human intervention is only expected in the event of an emergency (Jegadeesan and Venkatesan, 2016). Although such a model system is expected to be a norm in the near future, steps are underway to implement these integrated systems at least partly in livestock and agriculture.

Contagious livestock diseases can result in economic losses and decreased productivity for cattle farms. Park *et al*. have developed a livestock monitoring system (LMS) integrated with a wireless sensor network to collect data on heart rate, breathing rate and cattle movement. The authors report an average correlation coefficient of 0.97 for the collected data. Such systems have the ability to change the landscape of animal farming by providing comprehensive information on animal wellbeing. These systems are also able to rapidly identify livestock diseases and prevent economic losses stemming from loss of productivity (Park and Ha, 2015).

**Figure 5.**
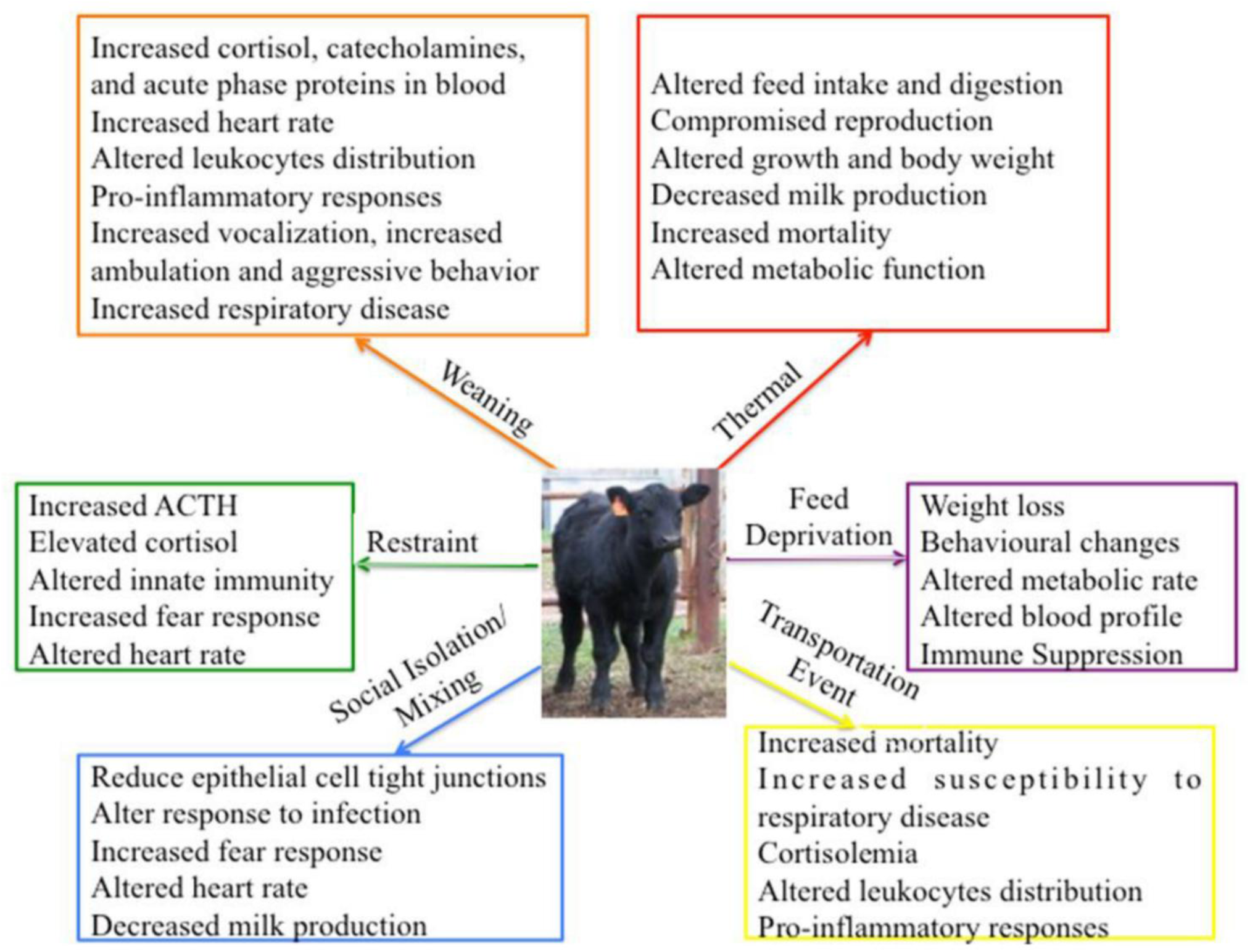
Impact of individual stressors on biological functions in cattle. Source: (Chen et al., 2015b)

### 9.1 Monitoring CO_2_ ventilation in farms

Carbon dioxide (CO_2_) can be used as a tracer gas to measure ventilation, as well as emission rates of CO_2_ (Persily, 2016). Currently, CO_2_ concentrations from agricultural facilities are measured using a Photo Acoustic Spectroscope (PAS) gas analyser (Chepete et al., 2012; Hassouna et al., 2013; Wheeler et al., 2006; Zhao et al., 2012) and the Open-Path laser (OP-laser) (Frish, 2014; He et al., 2009). Although these are appropriate for use in mechanically ventilated farms, the measurements in naturally ventilated farms show considerable variability. Moreover, these systems are cost-intensive when multiple samples are required. Non-Dispersive Infra-Red (NDIR) -based sensors are suggested as an alternative for PAS and OP-laser to measure CO_2_ concentrations in NV buildings. Experimental evaluation of NDIR sensors to measure CO_2_ levels has been demonstrated (Calvet et al., 2014; Piccot et al., 1994; Yasuda et al., 2012) along with the miniaturisation of the method, facilitating in-field usage (Hodgkinson et al., 2013). Mendes *et al*. have recently demonstrated the use of NDIR CO_2_ sensors for monitoring CO2 levels in a naturally ventilated dairy cow barn, comparing them to other commercially available NDIR CO_2_ sensors. The authors conclude that the CO_2_ concentrations were in agreement with the platforms tested, and the number of NDIR sensors required to represent the overall CO_2_ concentration of the dairy cow barn must be calculated based on the barn length and occupied barn area. The NDIR CO_2_ sensors were found suitable to be used as a multi-point monitoring system of CO_2_ concentrations in NV buildings as a feasible alternative to PAS and the OP-laser methods (Mendes et al., 2015).

### 9.2 Monitoring animal movement and behaviour

Monitoring movement and behaviour can provide information on an animal’s activity and wellbeing. A top-view camera can provide vital information if the animal is low-weight. Motion-detection technology and video recording coupled with the Gaussian Mixture Model (GMM) can be used to gather information on animal size and identify low-weight animals (Sa et al., 2015). MooMonitor integrates information on cow oestrus, as well as data on rumination, feeding and levels of activity. It makes use of wireless sensors for the two-way transmission of data. Other technologies, such as HerdNavigator™ and the Afimilk Silent Herdsman also serve the same purpose. The Silent Herdsman is a wearable technology and monitors all activities of cattle to analyse their behaviour. Any changes in an animal’s behaviour pattern can be used to identify the oestrus cycles and onset of disease/sickness.

Honeybees produce a variety of different sounds as a means to communicate with the colony. The sounds have characteristic low fundamental frequencies between 300 and 600 Hz (Barth et al., 2005). Honeybees’ sounds have specific frequencies within this range for a number of reasons. Both the range of sound frequency and the acoustic signal pattern determine the meaning of the sound. An accurate quantification of these signal patterns can give valuable information on the hive health (Qandour et al., 2014). Electronic systems for management of beehives have been developed lately and combine hive acoustics monitoring with measurement of parameters like brood temperature (Kridi et al., 2016), humidity, hive weight and the weather conditions of the apiary. Dietlein *et al*. have developed a system for automated continuous recording of sound emission by honeybees as a measure of their activity (Dietlein, 1985). The specific sounds from the hives are useful in providing information on colony behaviour, strength and health, and the data from the monitoring device can be accessed remotely from any internet-enabled device (Bromenshenk et al., 2015; Evans, 2015; Meikle and Holst, 2015). Other systems are being developed that integrate visual, acoustic and beehive monitoring systems and share them with the environmental monitoring platform (Figure 6). These systems can help in collecting and analysing the data on bee behaviour for biologists (Chiron et al., 2013).

**Figure 6.**
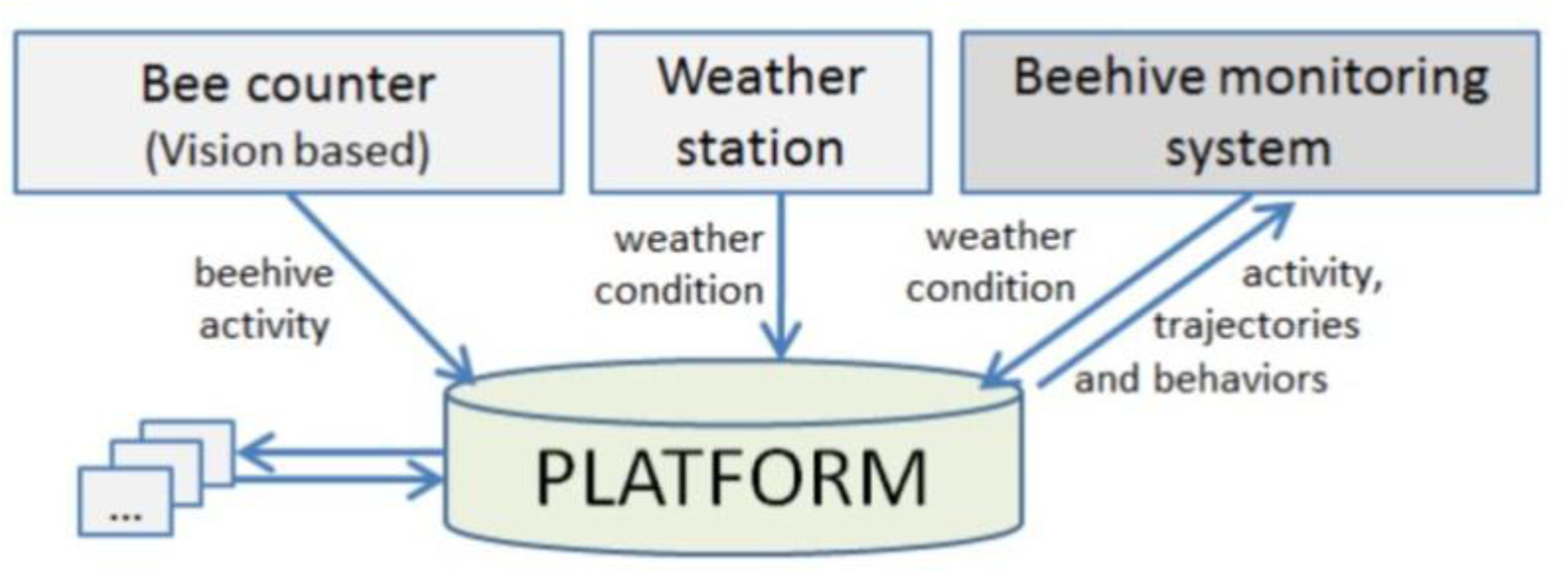
. Platform for gathering information from different blocks on bee activity, bee trajectories and weather. The platform also allows for sharing of information with the beehive monitoring system. Source: (Chiron et al., 2013).

### 9.3 Bioacoustic monitoring of poultry using biosensors

Livestock farming and production do not simply target economic goals, but food quality, safety, broiler production efficiency and sustainability (Berckmans, 2006). For these purposes, a growing need emerges in livestock farming, particularly in chicken farming, to monitor and assess the animals’ health, activity and welfare in real time, efficiently and economically.

Current, traditional monitoring systems in poultry farming are based on manual methods or simple systems relying on the observation, judgment and experience of the farmers, which is time-consuming, not real-time, and inaccurate.

Secondly, when it comes to the monitoring of the health, activity and welfare of chickens, currently available methods and systems cannot meet the increasing technical, administrative, and organisational requirements of ever-growing farms, which limits the possibility and feasibility of monitoring their livestock (Halachmi and Guarino, 2016). Thirdly, when targeting health and disease monitoring, clinical signs such as nasal discharge and diarrhoea are non-specific and cannot be used as evidence in diagnosis (Rahimian et al., 2012); and advanced diagnostic methods including ELISA and real-time RT-PCR are only useful when daily monitoring shows the necessity, because of their requirements for expert personnel, expensive equipment, and time-consuming and costly processes (Soltan et al., 2016).

Precision livestock farming and smart agriculture are calling for novel and automatic systems that are effective, efficient and affordable, and which meet variable goals to satisfy the purposes of monitoring the health, activity and welfare of chickens (Banakar et al., 2016). The ideal system would be an automatic device with an alarm feature, which is technologically simplified enough to be understandable and usable by farmers to assist in their daily monitoring of welfare, such as activities and conditions of the birds

Recent development of scientific research and technology in chickens and other livestock indicates that vocalisation monitoring could be a valuable tool for predicting diseases and enhancing productivity. Vocalisation technology is based on sounds made by the birds in their daily activities on the farms. The acoustic monitoring system was first developed to monitor coughing in porcine livestock farms, which found coughing to be an indicator of animal conditions, as it is a frequent symptom of multiple respiratory diseases affecting the lungs of livestock (Chedad et al., 2001). The new acoustic technology using neural networks as classification methods can distinguish cough sounds from other sounds such as metal clanging, grunts, and background noises. Using the nanobiosensors, vocalisation detection can be designed and developed as an efficient, effective and affordable system in monitoring the health conditions of chickens, to provide opportunities for making decisions and undertaking necessary actions.

Further development of the acoustic technology to detect porcine coughing contributed to a user-friendly computer software system called Cool Edit Program (Gutierrez et al., 2010). With the PC software system installed in porcine livestock farms, the farmers can monitor and initially diagnose pigs with wasting diseases on breeding farms in real time based on the online cough counter algorithm. An efficient data-mining technology was used to detect and recognise sounds from pigs with or without wasting diseases, and the sound data were collected in audio surveillance systems (Chung et al., 2013). Post sound acquisition, the widely used Mel Frequency Cepstrum Coefficient (MFCC) sound analysis technology was used to extract and differentiate sound data in the data pre-processing phase. Further development of the coughing sound recognition system used the Support Vector Data Description (SVDD) technology to enhance the automatic monitoring system. The SVDD worked as an anomaly or novelty detector to detect sound data related to porcine wasting diseases. Finally, the Sparse Representation Classifier (SRC) technology was used to classify the sound data based on their relations to different sub-types of porcine wasting diseases. Using the combination of MFCC, SVDD and SRC technology, the study yielded up to 94% disease detection and 91% classification accuracy of porcine wasting diseases.

The vocalisation- and acoustic-detection technology developed for pig coughing systems cannot be directly applied to poultry farms due to the types and the frequencies of the sounds the chickens make. The early exploration of the relation between vocalisation and poultry livestock welfare can be traced back to 1953 (Collias and Joos, 1953), and a great number of different vocalisation sounds, up to 30, have been described for juvenile and adult chickens (Manteuffel et al., 2004). The vocalisation system can allow the detection of signs of welfare as gakel-calls, alarm calls, distress calls, and stress calls through their sound analysis, indicating the levels of energy, frequency, and call duration, to detect normal welfare or impaired welfare. The socially relevant utterances will depend on complex contexts, but not solely on reactions to simple internal states and external stimuli. For example, the hens’ calls can be differentiated from cocks’ calls, and food calls from brooding hens to their chicks have a higher probability of occurrence when the chicks are not feeding or are at a distance (Wauters and Richard-Yris, 2002).

Bioacoustics from poultry farms can also help to identify genetic strains and sexes of the birds, in which the genetic strains can be identified using the second formant frequency and the pitch of the sounds; and the sexes can be identified using the second formant frequency apart from the sounds (Pereira et al., 2015a). An in-depth and comprehensive development of the acoustic system can detect the key vocalisation frequency and pattern changes in routine activities of the birds, and identify the age and weight of young broiler chickens based on the audio monitoring and comparison of anticipated sound patterns (Fontana et al., 2016). Figure 7 shows a schematic representation of how a well-designed intelligent technology using a vocalisation-detection system can monitor welfare and diagnose diverse infectious diseases in farmed chickens. Sound signals collected using microphones and a data collection card analysed by a neural network pattern-recognition system can detect and diagnose necrotic enteritis derived from the infection of *Clostridium perfringens* type A (Sadeghi et al., 2015). The diagnostic accuracy was 66.6% on day 2, and 100% on day 8 post the disease onset. Another study compared three different sound-detecting systems in the diagnosis of three avian infectious diseases with heavy economic losses: Newcastle Disease, Bronchitis Virus, and Avian Influenza (Banakar et al., 2016). The three systems were sound and time frequency detection alone, the Support Vector Data Description system (SVDD) (Chung et al., 2013), and an artificially intelligent device using SVDD as an input for intelligent analysis (Sadeghi et al., 2015). The results showed that the diagnostic accuracy from frequency detection was 41.4%, from SVDD was 83.3%, and from the artificially intelligent device was 91.2%. These results indicate that the combination of SVDD technology and artificially intelligent technology could yield the most accurate diagnosis in detecting poultry diseases from bioacoustics.

**Figure 7.**
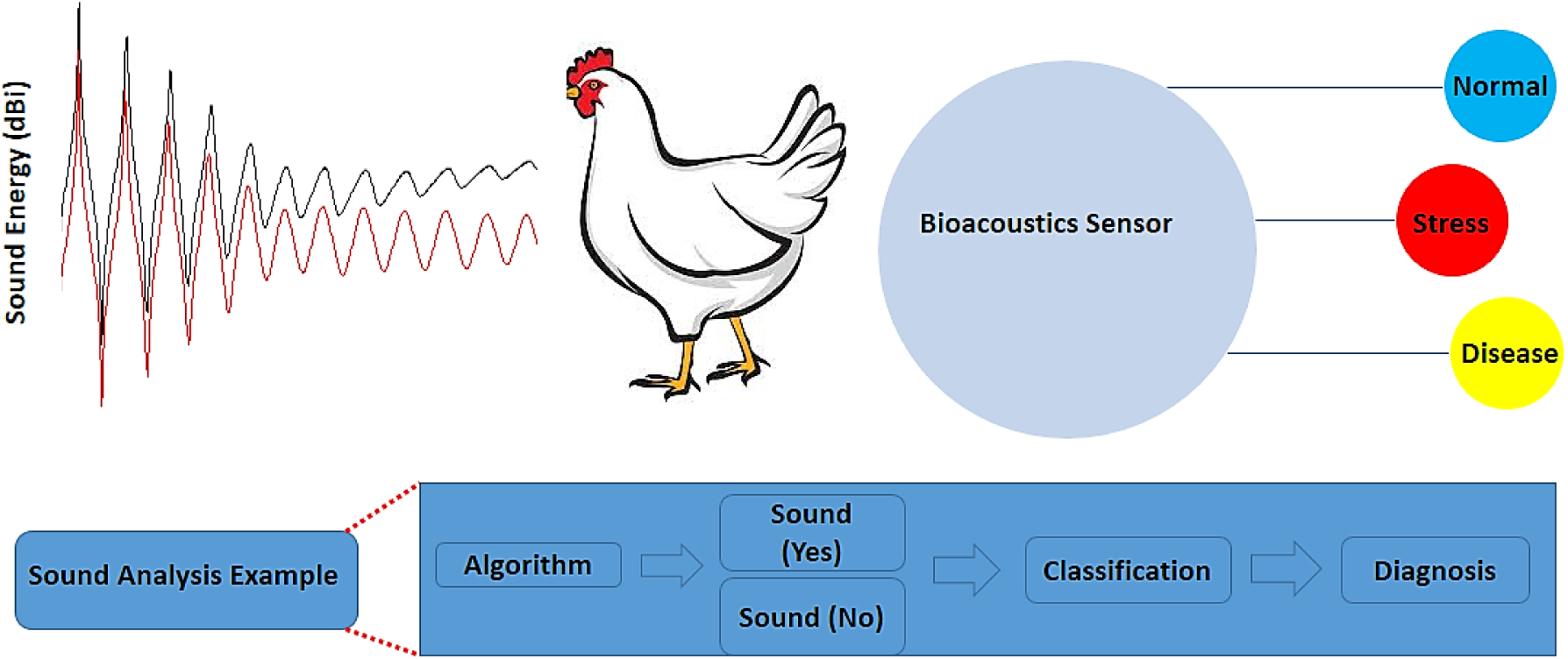
.Schematic representation of the use of bioacoustics analysis for detection of wellbeing in poultry livestock.

## 10. Perspectives

Precision livestock farming aims at creating a management system that relies upon autonomous, continuous, real-time monitoring and control of all aspects of livestock management, including reproduction, animal health and welfare, and the environmental impact of livestock production. It is assumed that the direct monitoring of animals will achieve greater control over their health status, which will eventually translate into better animal product quality over longer periods of time. Biosensor technology shall enable accurate and affordable acquisition of data points, while the smart algorithms, coupled with networked farms, shall further decision making and management processes in the animal farms. The primary goal of precision livestock farming is to generate reliable data using biosensors and run it through intelligent software systems to create value for the farmer, the environment, and the animals in the form of improved animal health and welfare, increased productivity and yields and reduced costs while minimising the impact on the environment.

While the biosensor technology is available for individual parameters, key advancements in the field are expected to generate robust monitoring systems for a multitude of parameters. Another key challenge currently faced is the slow uptake of these technologies on commercial farms. This has been attributed to the fact that although the precision systems and biosensors generate abundant data, the data is currently not being converted into useful information that could be utilised for the decision-making process in livestock management. Furthermore, the economic benefits of using these advanced systems is set to be demonstrated to individual farmers, who are reluctant to make investments in these systems in the absence of a clear economic benefit.

There is no doubt that advancements in the development of nanobiosensors, combining nanotechnology with highly specific analytic techniques for metabolic biomolecules and surveillance systems for monitoring animal health and welfare will be ubiquitously used to manage livestock farms and prevent disease outbreak. The key challenges that remain to be resolved include harmonisation of methods across various platforms and large-scale implementation of data analysis and sharing technologies.

## 11. Acknowledgments

The authors sincerely thank the Natural Sciences and Engineering Research Council of Canada (400929) and the Ontario Ministry of Agriculture, Food and Rural Affairs (300512) for funding this study.

## Highlights

- Advances in the signaling strategies and data acquisition for animal health management are summarized.
- Review focuses on the systems for observing physiological parameters and health of livestock.
- Nanomaterials based development and application of biosensors for animal health are discussed.
- Advancement in non-invasive biosensing techniques for the monitoring of livestock are reviewed.

## 12.References

Alexandratos, N., Bruinsma, J., 2012. World agriculture towards 2030/2050: the 2012 revision. ESA Working paper Rome, FAO.

Andersson, L.M., Okada, H., Miura, R., Zhang, Y., Yoshioka, K., Aso, H., Itoh, T., 2016. Wearable wireless estrus detection sensor for cows. Computers and Electronics in Agriculture 127, 101–108.

Andersson, L.M., Okada, H., Zhang, Y., Itoh, T., Miura, R., Yoshioka, K., 2015. Wearable wireless sensor for estrus detection in cows by conductivity and temperature measurements. IEEE SENSORS, pp. 1–4.

Andriamandroso, A., Lebeau, F., Bindelle, J., 2015. Changes in biting characteristics recorded using the inertial measurement unit of a smartphone reflect differences in sward attributes. In: M., G., D., B. (Eds.), Proceedings of the 7th European conference on Precision Livestock Farming, Precision Livestock Farming, pp. 283–289, Milan, Italy.

Bai, H., Wang, R., Hargis, B., Lu, H., Li, Y., 2012. A SPR Aptasensor for Detection of Avian Influenza Virus H5N1. Sensors 12(9), 12506–12518.

Banakar, A., Sadeghi, M., Shushtari, A., 2016. An intelligent device for diagnosing avian diseases. Comput. Electron. Agric. 127, 744–753.

Bandodkar, A.J., Wang, J., 2014. Non-invasive wearable electrochemical sensors: a review. Trends in Biotechnology 32(7), 363–371.

Barth, F.G., Hrncir, M., Tautz, J., 2005. Vibratory and airborne-sound signals in bee communication (Hymenoptera). Insect Sounds and Communication: Physiology, Behaviour, Ecology, and Evolution, pp. 421–436. CRC Press.

Benvenutti, M.A., Pavetti, D.R., Poppi, D.P., Gordon, I.J., Cangiano, C.A., 2016. Defoliation patterns and their implications for the management of vegetative tropical pastures to control intake and diet quality by cattle. Grass and Forage Science 71(3), 424–436.

Berckmans, D., 2006. Automatic on-line monitoring of animals by precision livestock farming. Livestock production and society 287.

Bhatta, D., Villalba, M.M., Johnson, C.L., Emmerson, G.D., Ferris, N.P., King, D.P., Lowe, R., 2012. Rapid Detection of Foot-and-Mouth Disease Virus with Optical Microchip Sensors. Procedia Chemistry 6, 2–10.

Blom, J., Ridder, C., 2010. Reproductive Management and Performance Can be Improved by Use of DeLaval Herd Navigator®. The First North American Conference on Precision Dairy Management.

Braun, U., Trösch, L., Nydegger, F., Hässig, M., 2013. Evaluation of eating and rumination behaviour in cows using a noseband pressure sensor. BMC Veterinary Research 9(1), 164.

Bromenshenk, J.J., Henderson, C.B., Seccomb, R.A., Welch, P.M., Debnam, S.E., Firth, D.R., 2015. Bees as biosensors: chemosensory ability, honey bee monitoring systems, and emergent sensor technologies derived from the pollinator syndrome. Biosensors 5(4), 678–711.

Burciaga-Robles, L.O., Holland, B.P., Step, D.L., Krehbiel, C.R., McMillen, G.L., Richards, J., Sims, L.E., Jeffers, J.D., Namjou, K., McCann, P.J., 2009. Evaluation of breath biomarkers and serum haptoglobin concentration for diagnosis of bovine respiratory disease in heifers newly arrived at a feedlot. American journal of veterinary research 70(10), 1291–1298.

Caja, G., Castro-Costa, A., Knight, C.H., 2016. Engineering to support wellbeing of dairy animals. Journal of Dairy Research 83(2), 136–147.

Calvet, S., Campelo, J.C., Estellés, F., Perles, A., Mercado, R., Serrano, J.J., 2014. Suitability evaluation of multipoint simultaneous CO2 sampling wireless sensors for livestock buildings. Sensors 14(6), 10479–10496.

Chedad, A., Moshou, D., Aerts, J.-M., Van Hirtum, A., Ramon, H., Berckmans, D., 2001. AP— animal production technology: recognition system for pig cough based on probabilistic neural networks. Journal of agricultural engineering research 79(4), 449–457.

Chen, L., Ye, S., Cai, K., Zhang, C., Zhou, G., He, Z., Han, H., 2015a. An aqueous platinum nanotube based fluorescent immuno-assay for porcine reproductive and respiratory syndrome virus detection. Talanta 144, 324–328.

Chen, Y., Arsenault, R., Napper, S., Griebel, P., 2015b. Models and Methods to Investigate Acute Stress Responses in Cattle. Animals 5(4), 0411.

Chen, Y., Huang, C.-H., Hou, C., Huo, D., Jin, G., 2013. Rapid and Label-Free Detection of Porcine Reproductive and Respiratory Syndrome Virus on Nanoscale by Biosensor Based on Imaging Ellipsometry. Integrated Ferroelectrics 145(1), 122–129.

Chepete, H., Xin, H., Mendes, L., Li, H., Bailey, T., 2012. Ammonia emission and performance of laying hens as affected by different dosages of Yucca schidigera in the diet. The Journal of Applied Poultry Research 21(3), 522–530.

Chiron, G., Gomez-Krämer, P., Ménard, M., 2013. Outdoor 3D Acquisition System for Small and Fast Targets. Application to honeybee monitoring at the beehive entrance. GEODIFF 2013, pp. 10–19, Barcelona, France.

Christensen, L.S., Brehm, K.E., Skov, J., Harlow, K.W., Christensen, J., Haas, B., 2011. Detection of foot-and-mouth disease virus in the breath of infected cattle using a hand-held device to collect aerosols. Journal of virological methods 177(1), 44–48.

Chung, Y., Oh, S., Lee, J., Park, D., Chang, H.-H., Kim, S., 2013. Automatic detection and recognition of pig wasting diseases using sound data in audio surveillance systems. Sensors 13(10), 12929–12942.

Clapham, W.M., Fedders, J.M., Beeman, K., Neel, J.P., 2011. Acoustic monitoring system to quantify ingestive behavior of free-grazing cattle. Computers and Electronics in Agriculture 76(1), 96–104.

Collias, N., Joos, M., 1953. The spectrographic analysis of sound signals of the domestic fowl. Behaviour 5(1), 175–188.

Cook, N.J., 2012. Review: Minimally invasive sampling media and the measurement of corticosteroids as biomarkers of stress in animals. Canadian Journal of Animal Science 92(3), 227–259.

Da Silva, N., Zardoya, R., Santurde, G., Solana, A., Castro, J., 1995. Rapid and sensitive detection of the bovine viral diarrhea virus genome in semen. Journal of virological methods 55(2), 209–218.

Davis, J.D., 2007a. Remote characterization of locomotion, grazing and drinking behavior in beef cattle using GPS and ruminant temperature dynamics. ProQuest.

Davis, J.D.L., 2007b. Remote Characterization of Locomotion, Grazing and Drinking Behavior in Beef Cattle Using GPS and Ruminant Temperature Dynamics. Iowa State University.

De Groot, D., Slieker, R., Van Loo, P., Verheij, E., Lorentsen, H., Laursen, M., Markert, M., Masse, F., Bouwens, F., Integrative Wireless Monitoring of Minipigs. Newsletter 38 Autumn 2012 pp 11–17.

Dietlein, D.G., 1985. A Method for Remote Monitoring of Activity of Honeybee Colonies by Sound Analysis. Journal of Apicultural Research 24(3), 176–183.

Diouani, M.F., Helali, S., Hafaid, I., Hassen, W.M., Snoussi, M.A., Ghram, A., Jaffrezic-Renault, N., Abdelghani, A., 2008. Miniaturized biosensor for avian influenza virus detection. Materials Science and Engineering: C 28(5–6), 580–583.

Duarte, C., Costa, T., Carneiro, C., Soares, R., Jitariu, A., Cardoso, S., Piedade, M., Bexiga, R., Freitas, P., 2016. Semi-Quantitative Method for Streptococci Magnetic Detection in Raw Milk. Biosensors 6(2), 19.

Durkin, J., DeLaval, B.W., 2010. Heat detection: Trends and opportunities. Proc. Second North Am. Conf. Precision Dairy Management, Toronto, Canada, pp. 1–10.

Ellis, C.K., Stahl, R.S., Nol, P., Waters, W.R., Palmer, M.V., Rhyan, J.C., VerCauteren, K.C., McCollum, M., Salman, M.D., 2014. A pilot study exploring the use of breath analysis to differentiate healthy cattle from cattle experimentally infected with Mycobacterium bovis. PLoS One 9(2), e89280.

Evans, S.K., 2015. Electronic beehive monitoring–applications to research. Julius-Kühn-Archiv 0(450), 121.

Fend, R., Geddes, R., Lesellier, S., Vordermeier, H.-M., Corner, L.A.L., Gormley, E., Costello, E., Hewinson, R.G., Marlin, D.J., Woodman, A.C., Chambers, M.A., 2005. Use of an Electronic Nose To Diagnose Mycobacterium bovis Infection in Badgers and Cattle. Journal of Clinical Microbiology 43(4), 1745–1751.

Fine, G.F., Cavanagh, L.M., Afonja, A., Binions, R., 2010. Metal oxide semi-conductor gas sensors in environmental monitoring. Sensors 10(6), 5469–5502.

Fontana, I., Tullo, E., Scrase, A., Butterworth, A., 2016. Vocalisation sound pattern identification in young broiler chickens. Animal: an international Journal of Animal Bioscience 10(09), 1567–1574.

Force, A. T., 2015. Critical role of animal science research in food security and sustainability. The National Academies Press, Washington, DC.

Frasconi, M., Mazzarino, M., Botre, F., Mazzei, F., 2009. Surface plasmon resonance immunosensor for cortisol and cortisone determination. Anal Bioanal Chem 394(8), 2151–2159.

Frish, M.B., 2014. Current and emerging laser sensors for greenhouse gas sensing and leak detection. SPIE Sensing Technology+ Applications, pp. 91010H‐‐91012. International Society for Optics and Photonics.

Fuentes, M., Tecles, F., Gutiérrez, A., Otal, J., Martínez-Subiela, S., Cerón, J.J., 2011. Validation of an Automated Method for Salivary Alpha-Amylase Measurements in Pigs (Sus Scrofa Domesticus) and its Application as a Stress Biomarker. Journal of Veterinary Diagnostic Investigation 23(2), 282–287.

Fűtő, P., Markus, G., Kiss, A., Adányi, N., 2012. Development of a Catalase-Based Amperometric Biosensor for the Determination of Increased Catalase Content in Milk Samples. Electroanalysis 24(1), 107–113.

Gajendragad, M.R., Kamath, K.N.Y., Anil, P.Y., Prabhudas, K., Natarajan, C., 2001. Development and standardization of a piezo electric immunobiosensor for foot and mouth disease virus typing. Veterinary Microbiology 78(4), 319–330.

Gao, W., Emaminejad, S., Nyein, H.Y.Y., Challa, S., Chen, K., Peck, A., Fahad, H.M., Ota, H., Shiraki, H., Kiriya, D., Lien, D.-H., Brooks, G.A., Davis, R.W., Javey, A., 2016. Fully integrated wearable sensor arrays for multiplexed in situ perspiration analysis. Nature 529(7587), 509–514.

Garner, C.E., Smith, S., Bardhan, P.K., Ratcliffe, N.M., Probert, C.S., 2009. A pilot study of faecal volatile organic compounds in faeces from cholera patients in Bangladesh to determine their utility in disease diagnosis. Transactions of the Royal Society of Tropical Medicine and Hygiene 103(11), 1171–1173.

Giovanetti, V., Decandia, M., Molle, G., Acciaro, M., Mameli, M., Cabiddu, A., Cossu, R., Serra, M.G., Manca, C., Rassu, S.P.G., Dimauro, C., 2017. Automatic classification system for grazing, ruminating and resting behaviour of dairy sheep using a tri-axial accelerometer. Livestock Science 196, 42–48.

Gomes, P., Giralt, E., Andreu, D., 1999. Surface plasmon resonance screening of synthetic peptides mimicking the immunodominant region of C-S8c1 foot-and-mouth disease virus. Vaccine 18(3-4), 362–370.

Grant, S.A., Heits, B., Kleiboeker, S., 2006. Development of an Optical Biosensor Utilizing GoldNanoparticles to Detect Porcine Reproductive and Respiratory Syndrome Virus. Sensor Letters 4(3), 246–252.

Gumus, A., Lee, S., Karlsson, K., Gabrielson, R., Winkler, D., Erickson, D., 2014. Real-time in vivo uric acid biosensor system for biophysical monitoring of birds. Analyst 139(4), 742–748.

Guo, D., Zhuo, M., Zhang, X., Xu, C., Jiang, J., Gao, F., Wan, Q., Li, Q., Wang, T., 2013. Indium-tin-oxide thin film transistor biosensors for label-free detection of avian influenza virus H5N1. Analytica Chimica Acta 773, 83–88.

Gutierrez, W., Kim, S., Kim, D., Yeon, S., Chang, H., 2010. Classification of porcine wasting diseases using sound analysis. Asian-Australasian Journal of Animal Sciences 23(8), 1096–1104.

Halachmi, I., Guarino, M., 2016. Editorial:Precision livestock farming: a ‘per animal’approach using advanced monitoring technologies. Animal: an international journal of animal bioscience 10(9), 1482–1483.

Hassouna, M., Robin, P., Charpiot, A., Edouard, N., Méda, B., 2013. Infrared photoacoustic spectroscopy in animal houses: Effect of non-compensated interferences on ammonia, nitrous oxide and methane air concentrations. Biosystems Engineering 114(3), 318–326.

He, Y., Zhang, Y.-j., Xia, H., Geng, H., Ruan, J., Wang, M., 2009. Open-path online monitoring of ambient atmospheric co2 based on laser absorption spectrum. Spectroscopy and Spectral Analysis 29(1), 10–13.

Hearps, A., Zhang, Z., Alexandersen, S., 2002. Evaluation of the portable Cepheid SmartCycler real-time PCR machine for the rapid diagnosis of foot-and-mouth disease. Veterinary Record 150(20), 625–628.

Heinze, B.C., Song, J.-Y., Lee, C.-H., Najam, A., Yoon, J.-Y., 2009. Microfluidic immunosensor for rapid and sensitive detection of bovine viral diarrhea virus. Sensors and Actuators B: Chemical 138(2), 491–496.

Hellsten, Y., Tullson, P.C., Richter, E.A., Bangsbo, J., 1997. Oxidation of urate in human skeletal muscle during exercise. Free radical biology & medicine 22(1-2), 169–174.

Herinaina, A.L., Bindelle, J., Mercatoris, B., Lebeau, F., 2016. A review on the use of sensors to monitor cattle jaw movements and behavior when grazing. Biotechnology, Agronomy, Society and Environment 23(S1), 273–286.

Hibi, K., Hatanaka, K., Takase, M., Ren, H., Endo, H., 2012. Wireless biosensor system for realtime L-lactic acid monitoring in fish. Sensors 12(5), 6269–6281.

Hodgkinson, J., Smith, R., Ho, W.O., Saffell, J.R., Tatam, R.P., 2013. Non-dispersive infra-red (NDIR) measurement of carbon dioxide at 4.2 μm in a compact and optically efficient sensor. Sensors and Actuators B: Chemical 186, 580–588.

Hogeveen, H., Kamphuis, C., Steeneveld, W., Mollenhorst, H., 2010. Sensors and Clinical Mastitis—The Quest for the Perfect Alert. Sensors (Basel, Switzerland) 10(9), 7991–8009.

Holtkamp, D.J., Kliebenstein, J.B., Neumann, E.J., 2013. Assessment of the economic impact of porcine reproductive and respiratory syndrome virus on United States pork producers. JSHAP 21(2), 72–84.

Iguchi, S., Kudo, H., Saito, T., Ogawa, M., Saito, H., Otsuka, K., Funakubo, A., Mitsubayashi, K., 2007. A flexible and wearable biosensor for tear glucose measurement. Biomedical Microdevices 9(4), 603–609.

Ingram, B., Gavine, F., Lawson, P., 2005. Fish Health Management Guidelines for Farmed Murray Cod., Fisheries Victoria Research Report.

Jegadeesan, S., Venkatesan, G.P., 2016. Smart cow health monitoring, farm environmental monitoring and control system using wireless sensor networks. Int. J. Adv. Eng. Tech./Vol VII/Issue I/Jan–March 334, 339.

Jia, W., Bandodkar, A.J., Valdés-Ramírez, G., Windmiller, J.R., Yang, Z., Ramírez, J., Chan, G., Wang, J., 2013. Electrochemical Tattoo Biosensors for Real-Time Noninvasive Lactate Monitoring in Human Perspiration. Analytical chemistry 85(14), 6553–6560.

Jindal, K., Tomar, M., Gupta, V., 2012. CuO thin film based uric acid biosensor with enhanced response characteristics. Biosensors and Bioelectronics 38(1), 11–18.

Jónsson, R., Blanke, M., Poulsen, N.K., Caponetti, F., Højsgaard, S., 2011. Oestrus detection in dairy cows from activity and lying data using on-line individual models. Computers and Electronics in Agriculture 76(1), 6–15.

Kim, J., Imani, S., de Araujo, W.R., Warchall, J., Valdés-Ramírez, G., Paixão, T.R.L.C., Mercier, P.P., Wang, J., 2015. Wearable salivary uric acid mouthguard biosensor with integrated wireless electronics. Biosensors and Bioelectronics 74, 1061–1068.

Kim, S.B., Takenaka, Y., Torimura, M., 2011. A Bioluminescent Probe for Salivary Cortisol. Bioconjugate Chemistry 22(9), 1835–1841.

Knobloch, H., Köhler, H., Commander, N., Reinhold, P., Turner, C., Chambers, M., Pardo, M., Sberveglieri, G., 2009. Volatile organic compound (VOC) analysis for disease detection: proof of principle for field studies detecting paratuberculosis and brucellosis. AIP Conference Proceedings, pp. 195–197. AIP.

Kridi, D.S., Carvalho, C.G.N.d., Gomes, D.G., 2016. Application of wireless sensor networks for beehive monitoring and in-hive thermal patterns detection. Comput. Electron. Agric.127, 221–235.

Kumanan, V., Nugen, S.R., Baeumner, A.J., Chang, Y.-F., 2009. A biosensor assay for the detection of Mycobacterium avium subsp. paratuberculosis in fecal samples. J Vet Sci 10(1), 35–42.

La Belle, J.T., Engelschall, E., Lan, K., Shah, P., Saez, N., Maxwell, S., Adamson, T., Abou-Eid, M., McAferty, K., Patel, D.R., Cook, C.B., 2014. A Disposable Tear Glucose Biosensor-Part 4: Preliminary Animal Model Study Assessing Efficacy, Safety, and Feasibility. J Diabetes Sci Technol 8(1), 109–116.

Laca, WallisDeVries, 2000. Acoustic measurement of intake and grazing behaviour of cattle. Grass and Forage Science 55(2), 97–104.

Lee, S.J., Kwon, Y.S., Lee, J.-e., Choi, E.-J., Lee, C.-H., Song, J.-Y., Gu, M.B., 2013. Detection of VR-2332 Strain of Porcine Reproductive and Respiratory Syndrome Virus Type II Using an Aptamer-Based Sandwich-Type Assay. Analytical chemistry 85(1), 66–74.

Leonardi, S., Marchesi, G., Tangorra, F.M., Lazzari, M., 2013. Use of a proactive herd management system in a dairy farm of northern italy: technical and economic results. Journal of Agricultural Engineering 44(2s).

Leopold, J.H., van Hooijdonk, R.T., Sterk, P.J., Abu-Hanna, A., Schultz, M.J., Bos, L.D., 2014. Glucose prediction by analysis of exhaled metabolites - a systematic review. BMC anesthesiology 14, 46.

Li, D., Wang, J., Wang, R., Li, Y., Abi-Ghanem, D., Berghman, L., Hargis, B., Lu, H., 2011. A nanobeads amplified QCM immunosensor for the detection of avian influenza virus H5N1. Biosensors and Bioelectronics 26(10), 4146–4154.

Liu, J., Chou, A., Rahmat, W., Paddon-Row, M.N., Gooding, J.J., 2005. Achieving direct electrical connection to glucose oxidase using aligned single walled carbon nanotube arrays. Electroanalysis 17(1), 38–46.

Lum, J., Wang, R., Lassiter, K., Srinivasan, B., Abi-Ghanem, D., Berghman, L., Hargis, B., Tung, S., Lu, H., Li, Y., 2012. Rapid detection of avian influenza H5N1 virus using impedance measurement of immuno-reaction coupled with RBC amplification. Biosensors and Bioelectronics 38(1), 67–73.

Luo, Y., Nartker, S., Miller, H., Hochhalter, D., Wiederoder, M., Wiederoder, S., Setterington, E., Drzal, L.T., Alocilja, E.C., 2010. Surface functionalization of electrospun nanofibers for detecting E. coli O157:H7 and BVDV cells in a direct-charge transfer biosensor. Biosensors and Bioelectronics 26(4), 1612–1617.

Malon, R.S.P., Sadir, S., Balakrishnan, M., #xf3, rcoles, E.P., 2014. Saliva-Based Biosensors: Noninvasive Monitoring Tool for Clinical Diagnostics. BioMed Research International 2014, 20.

Manteuffel, G., Puppe, B., Schön, P.C., 2004. Vocalization of farm animals as a measure of welfare. Applied Animal Behaviour Science 88(1), 163–182.

MarketsandMarkets, Precision Farming Market by Technology (Guidance System, Remote Sensing, Variable Rate Technology), Offering (Hardware Automation & Control System, Sensor & Monitoring Device, Software, Services), Application, and Geography - Global Forecast to 2022

Martinez-Pérez, D., Ferrer, M.L., Mateo, C.R., 2003. A reagent less fluorescent sol–gel biosensor for uric acid detection in biological fluids. Analytical Biochemistry 322(2), 238–242.

Mattachini, G., Riva, E., Perazzolo, F., Naldi, E., Provolo, G., 2016. Monitoring feeding behaviour of dairy cows using accelerometers. Journal of Agricultural Engineering 47(1), 54–58.

Mazeris, F., 2010. DeLaval herd navigator: proactive herd management. Proceedings of First North American Conference on Precision Dairy Management, pp. 26–27.

McNerney, R., Wondafrash, B.A., Amena, K., Tesfaye, A., McCash, E.M., Murray, N.J., 2010. Field test of a novel detection device for Mycobacterium tuberculosis antigen in cough. BMC infectious diseases 10, 161.

Meikle, W., Holst, N., 2015. Application of continuous monitoring of honeybee colonies. Apidologie 46(1), 10–22.

Mendes, L., Ogink, N., Edouard, N., van Dooren, H., Tinôco, I., Mosquera, J., 2015. NDIR Gas Sensor for Spatial Monitoring of Carbon Dioxide Concentrations in Naturally Ventilated Livestock Buildings. Sensors 15(5), 11239.

Milone, D.H., Galli, J.R., Cangiano, C.A., Rufiner, H.L., Laca, E.A., 2012. Automatic recognition of ingestive sounds of cattle based on hidden Markov models. Computers and electronics in agriculture 87, 51–55.

Mitchell, J., 2010. Small molecule immunosensing using surface plasmon resonance. Sensors (Basel, Switzerland) 10(8), 7323–7346.

Mitchell, J.S., Lowe, T.E., Ingram, J.R., 2009. Rapid ultrasensitive measurement of salivary cortisol using nano-linker chemistry coupled with surface plasmon resonance detection. Analyst 134(2), 380–386.

Mitchell, J.S., Wu, Y., Cook, C.J., Main, L., 2005. Sensitivity enhancement of surface plasmon resonance biosensing of small molecules. Anal Biochem 343(1), 125–135.

Montrose, A., Creedon, N., Sayers, R., Barry, S., O’riordan, A., 2015. Novel single gold nanowire-based electrochemical immunosensor for rapid detection of bovine viral diarrhoea antibodies in serum. Journal of Biosensors & Bioelectronics 6(3), 1–7.

Mottram, T., Berry, P., Pickard, A., Hart, J.P., Pemberton, R., 2004. Non-invasive system for monitoring physiological or health status of animals by sampling saliva. Google Patents.

Mottram, T., Dobbelaar, P., Schukken, Y., Hobbs, P., Bartlett, P., 1999. An experiment to determine the feasibility of automatically detecting hyperketonaemia in dairy cows. Livestock production science 61(1), 7–11.

Nadin, L.B., Chopa, F.S., Gibb, M.J., Trindade, J.K.d., Amaral, G.A.d., de Faccio Carvalho, P.C., Gonda, H.L., 2012. Comparison of methods to quantify the number of bites in calves grazing winter oats with different sward heights. Applied Animal Behaviour Science 139(1-2), 50–57.

Nakagawa, T., Hu, H., Zharikov, S., Tuttle, K.R., Short, R.A., Glushakova, O., Ouyang, X., Feig, I., Block, E.R., Herrera-Acosta, J., Patel, J.M., Johnson, R.J., 2006. A causal role for uric acid in fructose-induced metabolic syndrome. American journal of physiology. Renal physiology 290(3), F625–631.

Navon, S., Mizrach, A., Hetzroni, A., Ungar, E.D., 2013. Automatic recognition of jaw movements in free-ranging cattle, goats and sheep, using acoustic monitoring. Biosystems Engineering 114(4), 474–483.

Neethirajan, S., 2017. Recent advances in wearable sensors for animal health management. Sensing and Bio-Sensing Research 12, 15–29.

Neethirajan, S., Weng, X., Chen, L., 2016. Biosensor for detection of subclinical ketosis. Google Patents.

Neitzel, A.-C., Stamer, E., Junge, W., Thaller, G., 2014. Calibration of an automated California mastitis test with focus on the device-dependent variation. Springer Plus 3, 760.

Niedbalski, W., 2016. Recent progress in the diagnosis of foot-and-mouth disease: rapid field-based assays. Medycyna Weterynaryjna 72(6), 339–344.

Nydegger, F., Gyga, L., Egli, W., 2010. Automatic measurement of rumination and feeding activity using a pressure sensor. p. 027. Cemagref, Aubiere.

Nyhan, W.L., 1997. The recognition of Lesch-Nyhan syndrome as an inborn error of purine metabolism. Journal of inherited metabolic disease 20(2), 171–178.

Ospina, P., Nydam, D., Stokol, T., Overton, T., 2010. Associations of elevated nonesterified fatty acids and β-hydroxybutyrate concentrations with early lactation reproductive performance and milk production in transition dairy cattle in the northeastern United States. Journal of dairy science 93(4), 1596–1603.

Oudshoorn, F.W., Cornou, C., Hellwing, A.L.F., Hansen, H.H., Munksgaard, L., Lund, P., Kristensen, T., 2013. Estimation of grass intake on pasture for dairy cows using tightly and loosely mounted di- and tri-axial accelerometers combined with bite count. Comput. Electron. Agric. 99, 227–235.

Pahl, C., Hartung, E., Grothmann, A., Mahlkow-Nerge, K., Haeussermann, A., 2016. Suitability of feeding and chewing time for estimation of feed intake in dairy cows. Animal: an international journal of animal bioscience 10(9), 1507–1512.

Park, E.-J., Werner, J., Beebe, J., Chan, S., Barrie Smith, N., 2009. Noninvasive Ultrasonic Glucose Sensing with Large Pigs (∼200 Pounds) Using a Lightweight Cymbal Transducer Array and Biosensors. Journal of diabetes science and technology (Online) 3(3), 517–523.

Park, M.-C., Ha, O.-K., 2015. Development of effective cattle health monitoring system based on biosensors. Advanced Science and Technology Letters 117, 180–185.

Peled, N., Ionescu, R., Nol, P., Barash, O., McCollum, M., VerCauteren, K., Koslow, M., Stahl, R., Rhyan, J., Haick, H., 2012. Detection of volatile organic compounds in cattle naturally infected with Mycobacterium bovis. Sensors and Actuators B: Chemical 171-172, 588–594.

Pemberton, R.M., Hart, J.P., Mottram, T.T., 2001. An electrochemical immunosensor for milk progesterone using a continuous flow system1. Biosensors and Bioelectronics 16(9-12), 715–723.

Pereira, E.M., Naeaes, I.D.A., Garcia, R.G., 2015a. Vocalization of broilers can be used to identify their sex and genetic strain. Engenharia Agrícola 35(2), 192–196.

Pereira, J., Porto-Figueira, P., Cavaco, C., Taunk, K., Rapole, S., Dhakne, R., Nagarajaram, H., Camara, J.S., 2015b. Breath analysis as a potential and non-invasive frontier in disease diagnosis: an overview. Metabolites 5(1), 3–55.

Persily, A.K., 2016. Field measurement of ventilation rates. Indoor Air 26(1), 97–111.

Piccot, S.D., Masemore, S.S., Ringler, E.S., Srinivasan, S., Kirchgessner, D.A., Herget, W.F., 1994. Validation of a method for estimating pollution emission rates from area sources using open-path FTIR spectroscopy and dispersion modeling techniques. Air & Waste 44(3), 271–279.

Pritchard, G., Kirkwood, G., Sayers, A., 2002. Detecting antibodies to infectious bovine rhinotracheitis and BVD virus infections using milk samples from individual cows. Veterinary record 150(6), 182–183.

Qandour, A., Ahmad, I., Habibi, D., Leppard, M., 2014. Remote beehive monitoring using acoustic signals.

Rahimian, M., ZamaniI, M.A., Momtaz, H., Niazi, M., 2012. Detection and phylogenetic analysis of newcastle disease virus based on molecular techniques in broiler in Isfahan province.

Research, P.M., 2014. Biosensor Market Will Reach US $22,551.2 million in 2020. Persistence Market Research.

Reusch, C.E., Kley, S., Casella, M., 2006. Home monitoring of the diabetic cat. Journal of Feline Medicine & Surgery 8(2), 119–127.

Rose, D.P., Ratterman, M.E., Griffin, D.K., Hou, L., Kelley-Loughnane, N., Naik, R.R., Hagen, J.A., Papautsky, I., Heikenfeld, J.C., 2015. Adhesive RFID Sensor Patch for Monitoring of Sweat Electrolytes. IEEE Transactions on Biomedical Engineering 62(6), 1457–1465.

Ruiz-Garcia, L., Lunadei, L., Barreiro, P., Robla, I., 2009. A Review of Wireless Sensor Technologies and Applications in Agriculture and Food Industry: State of the Art and Current Trends. Sensors 9(6), 4728.

Rutter, S., Champion, R., Penning, P., 1997. An automatic system to record foraging behaviour in free-ranging ruminants. Applied Animal Behaviour Science 54(2-3), 185–195.

Rutter, S.M., 2000. Graze: A program to analyze recordings of the jaw movements of ruminants. Behavior Research Methods, Instruments, & Computers 32(1), 86–92.

Sa, J., Ju, M., Han, S., Kim, H., Chung, Y., Park, D., 2015. Detection of Low-Weight Pigs by Using a Top-View Camera. Proceedings of The fourth International Conference on Information Science and Cloud Computing (ISCC2015). 18–19 December 2015. Guangzhou, China. Online at http://pos.sissa.it/cgi-bin/reader/conf.cgi?confid=264,id.24.

Sadeghi, M., Banakar, A., Khazaee, M., Soleimani, M., 2015. An Intelligent Procedure for the Detection and Classification of Chickens Infected by Clostridium Perfringens Based on their Vocalization. Revista Brasileira de Ciência Avícola 17(4), 537–544.

Salómon, F., Tropea, S., Brengi, D., Hernández, A., Alamón, D., Parra, M., Longinotti, G., Ybarra, G., Lloret, P., Mass, M., Roberti, M., Lloret, M., Malatto, L., Moina, C., Fraigi, L., Melli, L., Cortina, M.E., Serantes, D.R., Ugalde, J.E., Ciocchini, A., Comerci, D.J., 2014.Smartphone controlled platform for point-of-care diagnosis of infectious diseases. 2014 IEEE 9th IberoAmerican Congress on Sensors, pp. 1–4.

Schaefer, A.L., Cook, N.J., Bench, C., Chabot, J.B., Colyn, J., Liu, T., Okine, E.K., Stewart, M., Webster, J.R., 2012. The non-invasive and automated detection of bovine respiratory disease onset in receiver calves using infrared thermography. Research in Veterinary Science 93(2), 928935.

Schazmann, B., Morris, D., Slater, C., Beirne, S., Fay, C., Reuveny, R., Moyna, N., Diamond, D., 2010. A wearable electrochemical sensor for the real-time measurement of sweat sodium concentration. Analytical Methods 2(4), 342–348.

Sethi, S., Nanda, R., Chakraborty, T., 2013. Clinical Application of Volatile Organic Compound Analysis for Detecting Infectious Diseases. Clinical microbiology reviews 26(3), 462–475.

Shao, K., Zhang, C., Ye, S., Cai, K., Wu, L., Wang, B., Zou, C., Lu, Z., Han, H., 2017. Near-infrared electrochemiluminesence biosensor for high sensitive detection of porcine reproductive and respiratory syndrome virus based on cyclodextrin-grafted porous Au/PtAu nanotube. Sensors and Actuators B: Chemical 240, 586–594.

Singh, A., Kaushik, A., Kumar, R., Nair, M., Bhansali, S., 2014. Electrochemical sensing of cortisol: a recent update. Applied biochemistry and biotechnology 174(3), 1115–1126.

Soltan, M.A., Tsai, Y.-L., Lee, P.-Y.A., Tsai, C.-F., Chang, H.-F.G., Wang, H.-T.T., Wilkes, R.P., 2016. Comparison of electron microscopy, ELISA, real time RT-PCR and insulated isothermal RT-PCR for the detection of Rotavirus group A (RVA) in feces of different animal species. Journal of virological methods 235, 99–104.

Soukup, M., Biesiada, I., Henderson, A., Idowu, B., Rodeback, D., Ridpath, L., Bridges, E.G., Nazar, A.M., Bridges, K.G., 2012. Salivary uric acid as a noninvasive biomarker of metabolic syndrome. Diabetology & metabolic syndrome 4(1), 14.

Spinhirne, J.P., Koziel, J.A., Chirase, N.K., 2004. Sampling and analysis of volatile organic compounds in bovine breath by solid-phase microextraction and gas chromatography-mass spectrometry. Journal of chromatography. A 1025(1), 63–69.

Stein, J.E., Greco, D.S., 2002. Portable blood glucose meters as a means of monitoring blood glucose concentrations in dogs and cats with diabetes mellitus. Clinical techniques in small animal practice 17(2), 70–72.

Stringer, R.C., Schommer, S., Hoehn, D., Grant, S.A., 2008. Development of an optical biosensor using gold nanoparticles and quantum dots for the detection of Porcine Reproductive and Respiratory Syndrome Virus. Sensors and Actuators B: Chemical 134(2), 427–431.

Su, X., Li, S.F.Y., Liu, W., Kwang, J., 2000. Piezoelectric quartz crystal based screening test for porcine reproductive and respiratory syndrome virus infection in pigs. Analyst 125(4), 725–730.

Sun, Y.-F., Liu, S.-B., Meng, F.-L., Liu, J.-Y., Jin, Z., Kong, L.-T., Liu, J.-H., 2012. Metal oxide nanostructures and their gas sensing properties: a review. Sensors 12(3), 2610–2631.

Tani, Y., Yokota, Y., Yayota, M., Ohtani, S., 2013. Automatic recognition and classification of cattle chewing activity by an acoustic monitoring method with a single-axis acceleration sensor. Computers and Electronics in Agriculture 92, 54–65.

Tarasov, A., Gray, D.W., Tsai, M.Y., Shields, N., Montrose, A., Creedon, N., Lovera, P., O’Riordan, A., Mooney, M.H., Vogel, E.M., 2016. A potentiometric biosensor for rapid on-site disease diagnostics. Biosensors & bioelectronics 79, 669–678.

Turner, C., Knobloch, H., Richards, J., Richards, P., Mottram, T.T.F., Marlin, D., Chambers, M.A., 2012. Development of a device for sampling cattle breath. Biosystems Engineering 112(2), 75–81.

Umemura, K., Wanaka, T., Ueno, T., 2009. Technical note: Estimation of feed intake while grazing using a wireless system requiring no halter. J Dairy Sci 92(3), 996–1000.

Ungar, E.D., Rutter, S.M., 2006. Classifying cattle jaw movements: comparing IGER behaviour recorder and acoustic techniques. Applied animal behaviour science 98(1), 11–27.

Vanrell, S.R., Chelotti, J.O., Galli, J., Rufiner, H.L., Milone, D.H., 2014. 3d acceleration for heat detection in dairy cows. XLIII Jornadas Argentinas de Informática e Investigación Operativa (43JAIIO)-VI Congreso Argentino de AgroInformática (CAI)(Buenos Aires, 2014).

Vreeburg, N., 2010a. Precision management on two Dutch dairy farms by use of Herd Navigator., First North American Conference on Precision Dairy Management, Toronto, Canada.

Vreeburg, N., 2010b. Precision Management On Two Dutch Dairy Farms By Use Of Herd Navigator®. First North American Conference on Precision Dairy Management, Toronto, Canada.

Wang, R., Li, Y., 2013. Hydrogel based QCM aptasensor for detection of avian influenzavirus. Biosensors and Bioelectronics 42, 148–155.

Wang, R., Wang, Y., Lassiter, K., Li, Y., Hargis, B., Tung, S., Berghman, L., Bottje, W., 2009. Interdigitated array microelectrode based impedance immunosensor for detection of avian influenza virus H5N1. Talanta 79(2), 159–164.

Waters, R.A., Fowler, V.L., Armson, B., Nelson, N., Gloster, J., Paton, D.J., King, D.P., 2014. Preliminary validation of direct detection of foot-and-mouth disease virus within clinical samples using reverse transcription loop-mediated isothermal amplification coupled with a simple lateral flow device for detection. PloS one 9(8), e105630.

Wathes, C.M., Kristensen, H.H., Aerts, J.M., Berckmans, D., 2008. Is precision livestock farming an engineer’s daydream or nightmare, an animal’s friend or foe, and a farmer’s panacea or pitfall? Computers and Electronics in Agriculture 64(1), 2–10.

Wauters, A.-M., Richard-Yris, M.-A., 2002. Mutual influence of the maternal hen’s food calling and feeding behavior on the behavior of her chicks. Developmental psychobiology 41(1), 25–36.

Weng, X., Chen, L., Neethirajan, S., Duffield, T., 2015a. Development of quantum dots-based biosensor towards on-farm detection of subclinical ketosis. Biosensors and Bioelectronics 72, 140–147.

Weng, X., Zhao, W., Neethirajan, S., Duffield, T., 2015b. Microfluidic biosensor for β-Hydroxybutyrate (βΗΒΑ) determination of subclinical ketosis diagnosis. Journal of Nanobiotechnology 13(1), 13.

Wheeler, E.F., Casey, K.D., Gates, R.S., Xin, H., Zajaczkowski, J.L., Topper, P.A., Liang, Y., Pescatore, A.J., 2006. Ammonia emissions from twelve US broiler chicken houses. Trans. ASABE 49(5), 1495–1512.

Wilson, A., 2015. Advances in Electronic-Nose Technologies for the Detection of Volatile Biomarker Metabolites in the Human Breath. Metabolites 5(1), 140.

Wu, H., Aoki, A., Arimoto, T., Nakano, T., Ohnuki, H., Murata, M., Ren, H., Endo, H., 2015. Fish stress become visible: A new attempt to use biosensor for real-time monitoring fish stress. Biosensors and Bioelectronics 67, 503–510.

Wu, S., Zhu, Y., Cai, Q., Zeng, K., Grimes, C.A., 2007. A wireless magnetoelastic α-amylase sensor. Sensors and Actuators B: Chemical 121(2), 476–481.

Xu, J., Suarez, D., Gottfried, D.S., 2007. Detection of avian influenza virus using an interferometric biosensor. Analytical and Bioanalytical Chemistry 389(4), 1193–1199.

Yamaguchi, M., Matsuda, Y., Sasaki, S., Sasaki, M., Kadoma, Y., Imai, Y., Niwa, D., Shetty, V., 2013. Immunosensor with fluid control mechanism for salivary cortisol analysis. Biosensors & bioelectronics 41, 186–191.

Yang, M., Caterer, N.R., Xu, W., Goolia, M., 2015. Development of a multiplex lateral flow strip test for foot-and-mouth disease virus detection using monoclonal antibodies. Journal of virological methods 221, 119–126.

Yang, M., Goolia, M., Xu, W., Bittner, H., Clavijo, A., 2013. Development of a quick and simple detection methodology for foot-and-mouth disease virus serotypes O, A and Asia 1 using a generic RapidAssay Device. Virology Journal 10, 125–125.

Yasuda, T., Yonemura, S., Tani, A., 2012. Comparison of the characteristics of small commercial NDIR CO2 sensor models and development of a portable CO2 measurement device. Sensors 12(3), 3641–3655.

Ye, W.W., Tsang, M.-K., Liu, X., Yang, M., Hao, J., 2014. Upconversion Luminescence Resonance Energy Transfer (LRET)-Based Biosensor for Rapid and Ultrasensitive Detection of Avian Influenza Virus H7 Subtype. Small 10(12), 2390–2397.

Yonemori, Y., Takahashi, E., Ren, H., Hayashi, T., Endo, H., 2009. Biosensor system for continuous glucose monitoring in fish. Analytica Chimica Acta 633(1), 90–96.

Zeidan, E., Shivaji, R., Henrich, V.C., Sandros, M.G., 2016. Nano-SPRi Aptasensor for the Detection of Progesterone in Buffer. Scientific Reports 6, 26714.

Zhao, Y., Pan, Y., Rutherford, J., Mitloehner, F.M., 2012. Estimation of the interference in multi-gas measurements using infrared photoacoustic analyzers. Atmosphere 3(2), 246–265.

